# Combining a homogenous KIM-1-DM1 antibody drug conjugate with sunitinib in renal cell carcinoma

**DOI:** 10.64898/2026.01.12.699160

**Authors:** Liya Thurakkal, Krishna Velayutham, Binayak Sarkar, Akshaya Kalyanaraman, Hae Lin Jang, Shiladitya Sengupta, Tanmoy Saha

## Abstract

Renal cell carcinoma (RCC) is the most common form of kidney cancer. It is also one of deadliest cancers, with a 5-year survival rate of less than 20% in advanced cancer patients with distant metastasis. Current treatments rely on targeted therapies, such as sunitinib, which are limited in eHicacy. Recently, antibody drug conjugates (ADCs) have emerged as a promising treatment modality by delivering cytotoxic payloads specifically to cancer cells. Here, we describe the engineering of a novel ADC (LT-025), where the antibody targets the kidney injury molecule (KIM) −1 receptor over-expressed on RCC cells to deliver a toxic maytansinoid (DM1) payload. Unlike prior attempts to engineer a KIM-1 targeted ADC that were limited by a heterogenous product, here we used a microbial transglutaminase (MTGase)-based conjugation strategy, which achieved a site-specific conjugation of the drug-linker to the antibody, resulting in a homogenous ADC with a drug-to-antibody ratio (DAR) of 2. The ADC exhibited prolonged stability, excellent antigen-binding capability, and KIM-1 expression-dependent cellular internalization and cytotoxicity in RCC, including in sunitinib-resistant RCC. Excitingly, LT-025 was well tolerated *in vivo*, and combining LT-025 and sunitinib exhibited a synergistic antitumor eHicacy in a RCC mouse model. Combining an ADC with a targeted therapeutic could emerge as a paradigm shift in the management of advanced RCC.

## INTRODUCTION

Kidney cancer is one of the ten most prevalent cancers worldwide with over 430,000 cases each year.^1^ Renal Cell Carcinoma (RCC) comprises nearly 90% of all kidney cancer types.^2^ The majority of the early-stage RCC patients are treated with surgery, but the overall outcome remains poor because of relapse and metastasis.^3,4^. Indeed, more than 35% of the RCC patients experience metastasis to distant organs, and in such advanced cancer patients the 5-year survival remains a dismal 10-15%.^5^ Targeted therapy, such as sunitinib, an oral tyrosine kinase inhibitor (TKI), is approved as a first line treatment for advanced/metastatic RCC and as adjuvant treatment for high-risk RCC following nephrectomy, but is limited by insufficient efficacy and adaptive treatment resistance.^6–8^ As a result, more than 170,000 RCC-related deaths occur worldwide. There is an unmet need to introduce an effective treatment strategy for RCC.

Over the last decade, the use of antibody-drug conjugates (ADCs) has gained attention because of their capability of delivering toxic payloads specifically to cancer cells.^9^ Antibodies against cancer cell-specific antigens are linked with a cytotoxic payload via a linker to achieve ADCs. Currently, several ADCs have shown promising outcomes in the clinic.^10,11^ We rationalized an ADC, especially in combination with a targeted therapeutic, could result in enhanced antitumor efficacy against RCC. Interestingly, an ADC, known as CDX-014, targeting kidney injury molecule-1 (KIM-1) was indeed in development for treating RCC, but it failed during clinical evaluation due to toxicity.^12,13^, potentially attributed to the premature release of the payload^14^. Indeed, off-target toxicity, premature release of payload, and the heterogeneous conjugation of the antibody and the drug are a few reasons for the failure of many ADCs in clinical trials.^15,16–19^ We rationalized that an ideal ADC for RCC needed to address these challenges.

To address the above challenges, here we engineered a homogenous KIM-1 targeted ADC, using a microbial transglutaminase (MTGase)-mediated transamidation reaction to conjugate a drug-linker payload to the Fc region of the antibody. KIM-1, a glycoprotein, is absent in normal kidney cells but is upregulated in kidney cancer, and hence is an ideal antigen to target.^20–23^ We used DM1 (mertansine) as the cytotoxic payload, which has been included in multiple clinically-used ADCs. We have extensively characterized this novel ADC, and tested it for efficacy and safety *in vivo*, in RCC. Our results show, combining the ADC with a targeted therapeutic, sunitinib, could shift the paradigm in the management of advanced RCC.

## RESULTS

### KIM-1 expression in renal cell carcinoma tissue and adjacent normal tissue

We first confirmed the KIM-1 overexpression in RCC tissues over other normal adjacent tissues. According to The Cancer Genome Atlas (TCGA) database, the median expression of KIM-1 transcript per million (TPM) in kidney cancer and normal kidney cells is 25.73 and 3.6, respectively (**Supplementary Fig.1a**). The Gene Expression Profiling Interactive Analysis (GEPIA) analysis of the KIM-1 revealed its expression exclusively in the kidney and expressively upregulated in the case of kidney cancer (**Fig.1a, supplementary Fig.1**). Next, we performed immunohistochemistry to investigate the expression of the KIM-1 in RCC patient-derived tumor samples together with the normal adjacent kidney tissue (NAT) from 16 RCC patients randomized based on age (26 to 79 years), sex, and cancer stage (I to III). Immunostaining of the tissue samples revealed high KIM-1 expression in RCC specimens (**Fig.1b**). We quantified the KIM-1 intensity and normalized it with the tissue density for each cancer and normal adjacent tissue (NAT). We found a significantly higher KIM-1 expression in cancer tissue than its matched NAT in 11 out of 16 patients (**Fig.1c**), consistent with the 70% KIM-1 positive RCC patients’ data in the literature.^20,21,23–26^ The significantly higher KIM-1 expression in cancer tissue in the majority of the RCC patients validates KIM-1 as a promising target for the antibody-drug conjugate.

**Figure 1:**
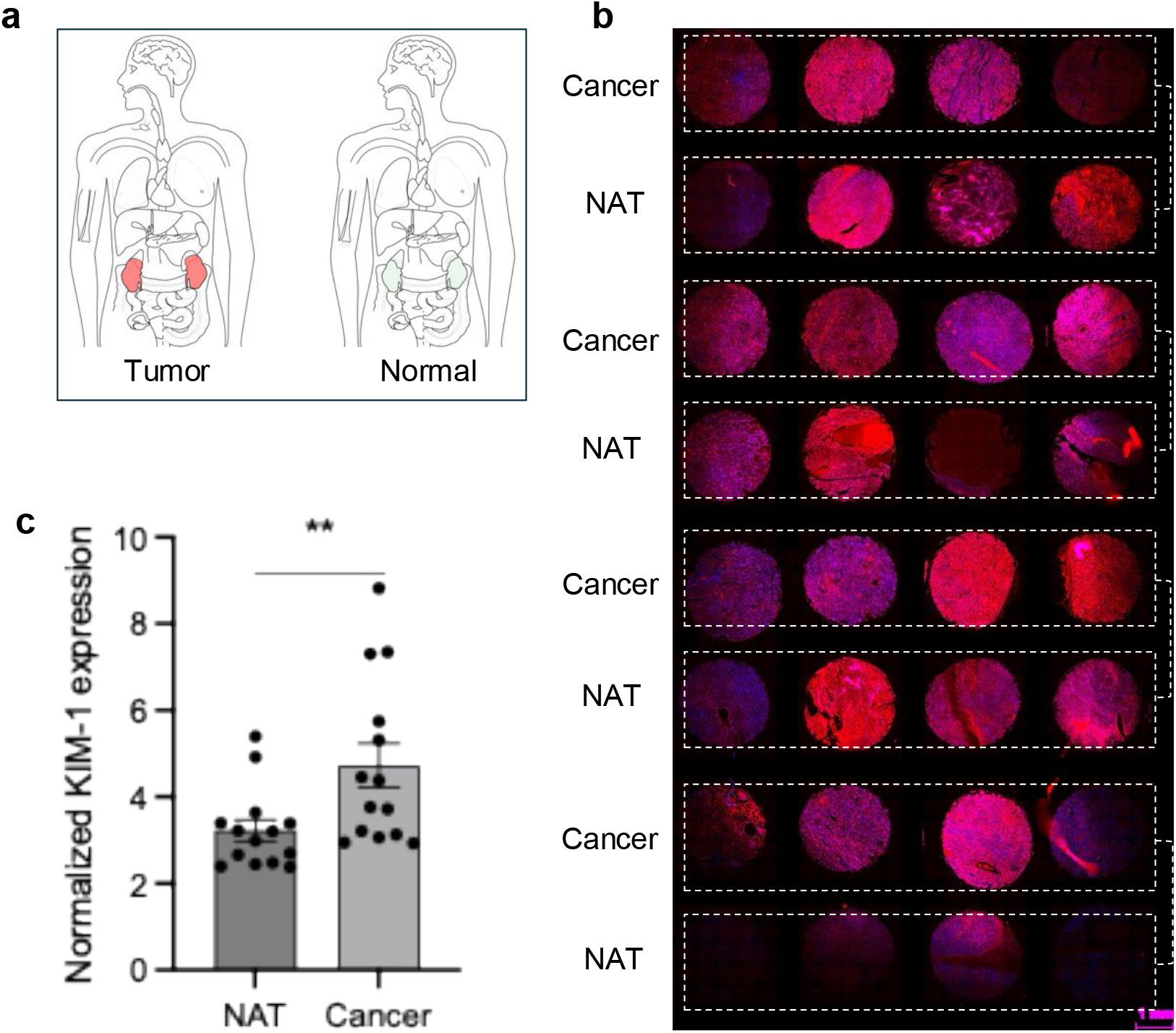
KIM-1 overexpression in RCC. (**a**) Median expression of the KIM-1 gene in different cancer and normal tissues obtained from the TCGA dataset using Gene Expression Profiling Interactive Analysis (GEPIA). The pictorial representation of the KIM-1 expression in cancer (red) and normal (green) organs. The figure was generated using the GEPIA platform. (**b**) Immunostaining of kidney tissue microarray by anti-KIM-1 (red) antibody, and DAPI (blue). Tissue specimens were collected from randomized patients based on age, sex, and stage of cancer. The normal adjacent tissue (NAT) and tumor tissue were collected from the same patients. The image shows the majority of the tumor expressing high KIM-1. 25 individual images were combined to form a single circular tissue image. (**c**) Bar plot showing relative KIM-1 expression in cancer and NAT. Each data point represents one patient sample. The data were represented as mean ± SEM (n = 16, paired t-test, P < 0.01 **).

### LT-025 (KIM-1 ADC) development and biophysical characterization

We selected an enzymatic site-specific conjugation of the drug-linker at the Fc region of the antibody to avoid any interference with the antigen-binding site. We used Mertansine (DM1), a potent microtubule inhibitor, as the payload. DM1 is a popular payload for ADC design because of its excellent cytotoxicity.^27,28^ At first, we synthesized the drug-linker, DM1-PEG_9_-NH_2_, by conjugating DM1 with a PEG_9_-NH_2_ linker using thiol maleimide chemistry (**supplementary Fig.2a**). We characterized the DM1-linker with ^1^H Nuclear Magnetic Resonance (NMR), ^13^C NMR, and mass spectrometry (**supplementary Fig.2b**,**c, and 3b**). The purity of the linker was determined using High Performance-Liquid Chromatography (HPLC) before proceeding with the antibody conjugation (**supplementary Fig.3a**).

We next used Microbial Transglutaminase-mediated (MTGase) conjugation, which exclusively conjugates the drug-linker at the Q295 position of CH2 domain of the Fc region of the IgG1 antibody.^29^ However, in the native form of the antibody, Q295 is not accessible to the MTGase because of the presence of N-glycans at the N297 location. Hence, we first deglycosylated the native antibody using a standard PNGase-F method, which selectively removes the N-linked oligosaccharide unit from the N297 position (**Fig.2a**).^30,31^ We monitored the deglycosylation reaction using hydrophobic interaction chromatography (HIC) and confirmed with electron spray ionization-mass spectrometry (ESI-MS) (**Fig.2b,c**). The HIC chromatogram showed an increase in retention time, consistent with the change in the hydrophobicity of the antibody (**Fig.2b**) upon removal of N-glycan. The observed loss of 1445 Da mass in ESI-MS of heavy chain confirmed the successful deglycosylation from heavy chain region with no change in the light chain mass (**Fig.2c**). Next, the DM1-PEG_9_-NH_2_ linker was attached to the antibody in the presence of MTGase, which catalyzes an acyl transfer reaction between the γ-carboxyamide group of a glutamine (Q295) on the antibody with a primary amine. We optimized the MTGase conjugation of the drug-linker with the deglycosylated antibody to yield a homogeneous ADC with maximum yield (**supplementary Fig.4**). The single peak at the HIC chromatogram with a higher retention time than the deglycosylated antibody confirms the exclusive formation of the desired ADC. An increase of 1327 Da mass of the heavy chain confirms the conjugation of a single drug-linker per heavy chain (**Fig.1b,c**). We observed increased yield of the ADC while using the Enhanced Microbial Transglutaminase (eMTGase) over the classical MTGase (Activa TI Transglutaminase). However, using eMTGase often results in a hyper-conjugated product with more than one drug conjugation in each heavy chain, as indicated in some of the previous reports.^32^ We performed detailed optimization of reaction conditions in order to achieve a homogeneous ADC. **Supplementary Fig.4a**,**b** represents the optimized reaction condition with a single ADC product. We found that the deglycosylation step is essential to achieve a homogeneous ADC with one conjugation per heavy chain of the antibody. To verify that we attempted the transamidation reaction with the native antibody without the deglycosylation step. This reaction resulted in most of the unreacted antibody with one minor product, which did not match our desired ADC (**Fig.2d**).

**Figure 2:**
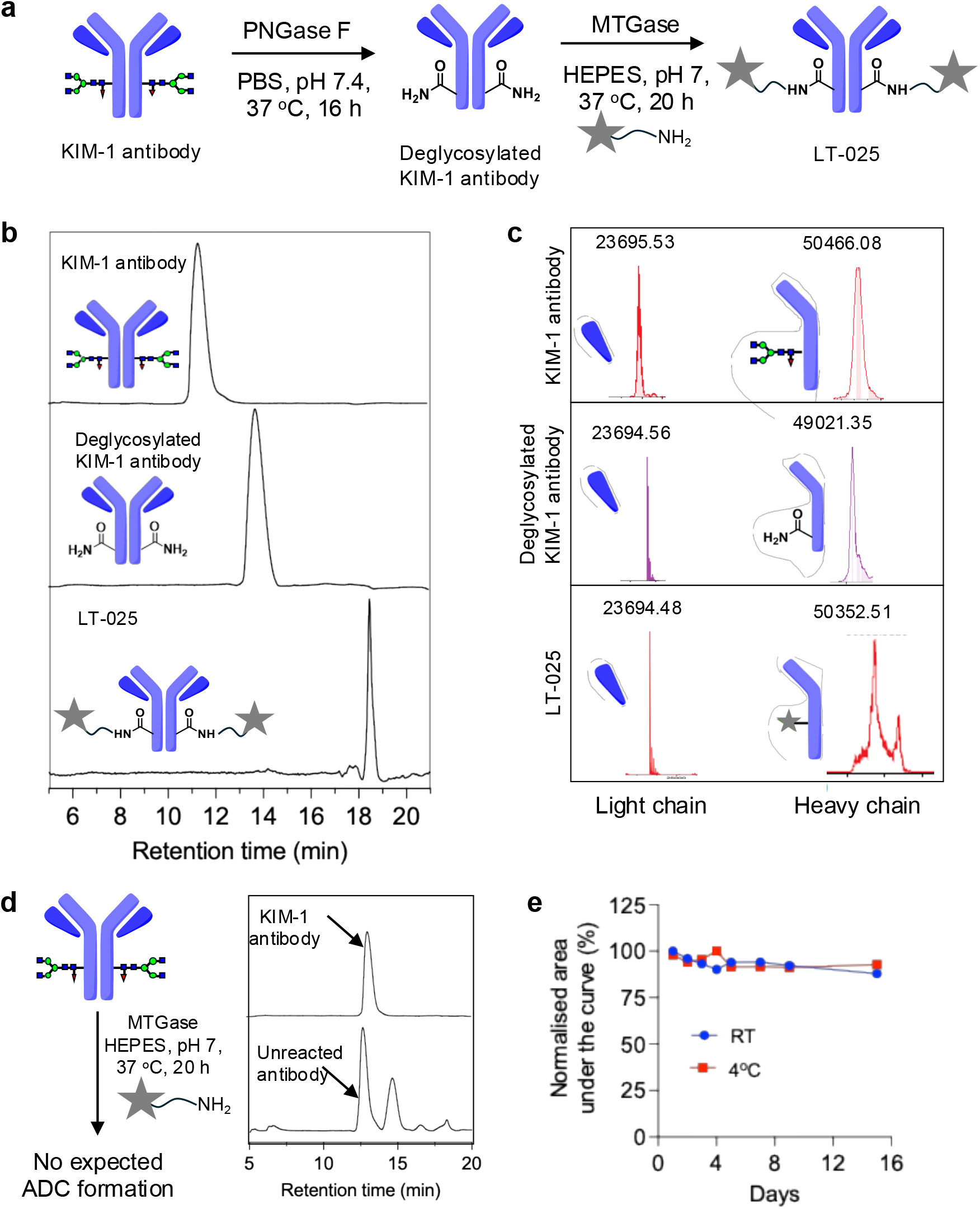
Preparation and characterization of KIM-1 targeting ADC, LT-025. (**a**) Schematic representation of the synthesis of KIM-1 targeting ADC, LT-025, by using transglutaminase. First, the native monoclonal KIM-1 antibody was deglycosylated in the presence of the PNGase F enzyme, and then the payload-linker was attached to the antibody at Q295 in the presence of the microbial transglutaminase enzyme (MTGase). The payload used was DM1. (**b**) Data shows the difference in the retention time of KIM-1 antibody, deglycosylated KIM-1 antibody, and ADC (LT-025) using hydrophobic interaction chromatography (HIC). The data shows single product formation in each step. Data was recorded under physiological conditions in PBS (pH 7.4) at 280 nm. 10 μg of each sample was injected for analysis. The chromatograms were baseline corrected using Origin 8.0 and plotted using GraphPad Prism 10 software. (**c**) Mass spectral analysis of the antibody drug conjugate. The data were recorded using ESI-MS after DTT reduction of the LT-025. Deconvoluted mass spectra represent the light chain and heavy chain of KIM-1 antibody, deglycosylated KIM-1 antibody, and LT-025. The data shows the change in mass of the heavy chain, signifying the selectivity of the reaction. 5 μg of each sample was injected for ESI-MS analysis. (**d**) The schematic shows the importance of the deglycosylation of the antibody for ADC production. The upper histogram represents the HIC chromatogram of the native antibody. The lower histogram is the product after the conjugation reaction without performing the deglycosylation step. The chromatogram shows the presence of unreacted antibody with some additional product, which does not match the desired LT-025 chromatogram. (**e**) The data represent the stability of the LT-025 for 15 days. The area under the curve from the HIC chromatogram of the ADC was normalized to day 1 and plotted with increasing days. 10 μL of 1 mg/mL of each sample was injected for analysis. The data shows no significant change in the peak area, indicating the stability of the ADC.

We calculated the drug-to-antibody ratio (DAR) using UV-visible spectroscopy and confirmed via ESI-MS. At first, we calculated the molar extinction coefficient of the DM1-linker and antibody at 254 and 280 nm (**supplementary Fig.5**). The absorption of the ADC was checked at 254 and 280 nm, and the DAR was calculated as described in previously reported literature^33,34^. We observed a DAR of 2 for LT-025. This was also validated from the HIC and ESI-MS data, where one drug-linker is conjugated with each heavy chain of the antibody. This supports our hypothesis on the use of the MTGase-mediated bioconjugation to produce homogenous ADC. We next incubated LT-025 in plasma at 4 °C and at room temperature, and investigated the stability using the HIC chromatography. The area of the peak corresponds to the ADC in the HIC-chromatogram, which was plotted in **Fig.2e**. The data show no significant change in the stability in either of the conditions for 15 days, suggesting that LT-025 is stable in plasma for over 15 days.

The conjugation of the drug-linker with the antibody may impact its binding to the KIM-1 protein in cancer cells. We used an enzyme-linked immunosorbent assay (ELISA) to investigate the antigen recognition capability of the LT-025 and compared it with the native KIM-1 antibody. We used a KIM-1 protein-coated ELISA plate to investigate the binding with the KIM-1 antibody and LT-025. We confirmed that the ADC can bind with the antigen in a concentration-dependent manner (**Fig.3a**). The antigen-binding of LT-025 obtained by the ELISA matched the native antibody, signifying no loss of antigen-binding property during the conjugation. Additionally, we used a competitive binding assay where the antigen binding of LT-025 was compared with a 1:1 mixture of native antibody and ADC. We observed a 50% reduction in the ADC binding with the antigen, as the remaining 50% has been occupied by the native antibody (**Fig.3b**). This confirms the comparable antigen-binding capability of the LT-025 and the native antibody.

**Figure 3.**
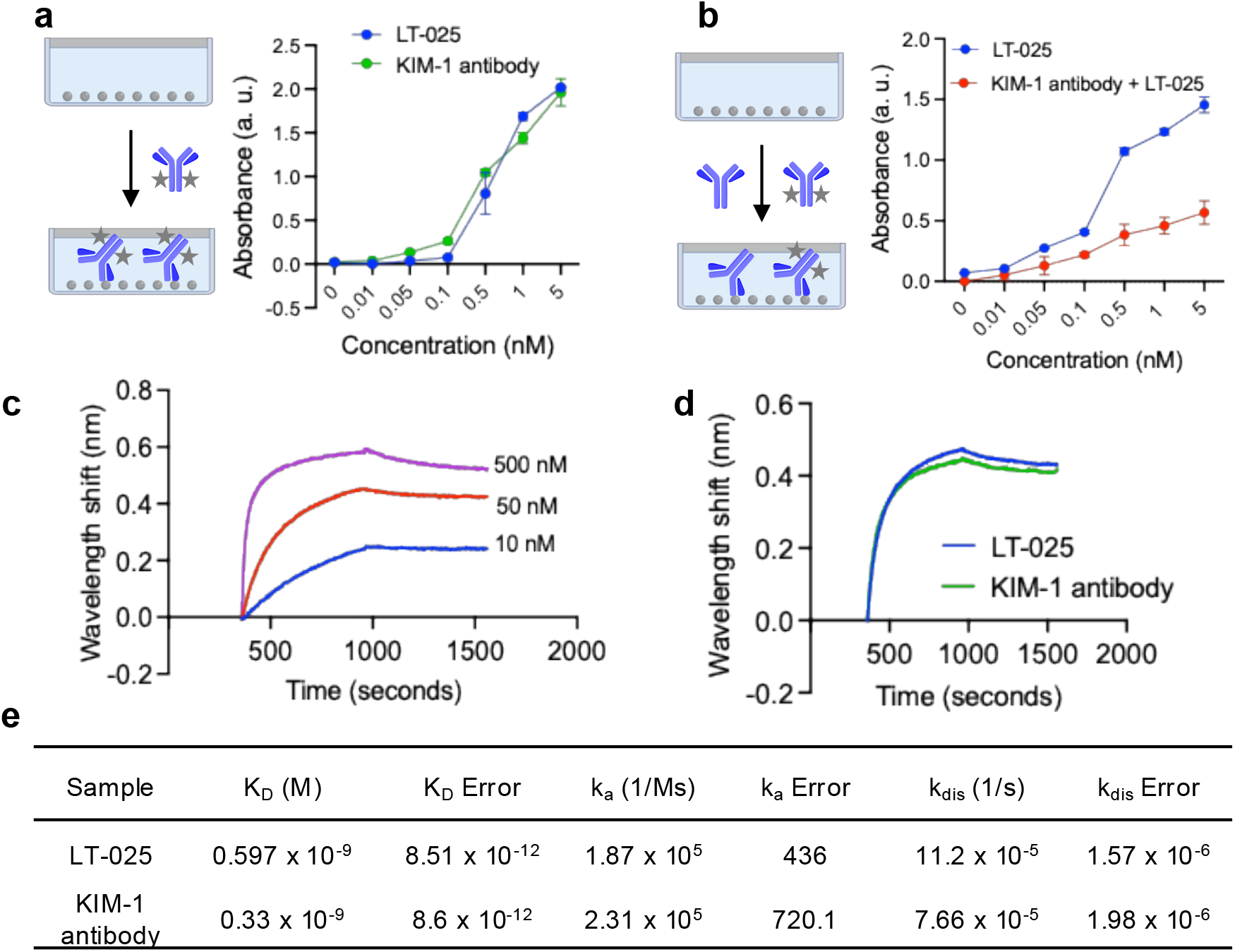
Antigen recognition by LT-025. (**a**) The antigen-binding capacity of LT-025 and the KIM-1 antibody was studied using ELISA. The immobilization of the LT-025 was checked in a KIM-1 protein-coated ELISA plate. The presence of the LT-025 was measured using an HRP-conjugated secondary antibody. The data show a dose-dependent increase in the absorption, representing the successful antigen binding by the LT-025. The antigen recognition of the LT-025 shows a similar dose response curve to that of the monoclonal KIM-1 antibody, signifying no loss of the antigen-binding capability of the ADC. (**b**) The comparison of the binding of LT-025 alone with ADC in the presence of KIM-1 free antibody was studied using ELISA. The data shows a 50% reduction in the immobilization of both ADC in the presence of free antibody. (**c**) Quantitative estimation of the antigen binding of the LT-025 using Bio-layer interferometry (BLI). The data show a dose-dependent increase in the association and dissociation events between LT-025 and KIM-1 protein-coated BLI sensor chip. (**d**) Comparison of the antigen binding of LT-025 and KIM-1 antibody using BLI, showing comparable antigen binding. 50 nM of each sample was used for the study. (**e**) Equilibrium constants and rate constants of LT-025 and the antibody (K_D_: Dissociation equilibrium constant, k_a_: association rate constant, and k_dis_: dissociation rate constant.

Further, to get quantitative insight into the antigen-binding ability of ADC, we used biolayer interferometry (BLI) analysis. The LT-025 showed a dose-dependent association and dissociation kinetics with the KIM-1 antigen-coated BLI sensor chip (**Fig.3c**). We observed a similar association and dissociation kinetics for LT-025 as the native KIM-1 antibody. The calculated dissociation constant of the LT-025 and the native antibody are *K*_D_ = 0.597 ± 0.008 nM, and *K*_D_ = 0.33 ± 0.008 nM, respectively (**Fig.3d**). This suggests that the deglycosylation followed by homogenous conjugation has no significant effect on antigen recognition as compared to the native antibody.

### KIM-1 dependent cellular internalization

Whether the antigen is internalized by the cell after antibody binding has implications on ADC design; ADCs that are internalized can leverage conjugation chemistries that are more stable as opposed to non-internalizing antigens, where the payload needs to be released outside the cells. To test whether KIM-1 engagement results in internalization of the construct, we prepared an antibody-fluorophore conjugate (AFC) as a surrogate for an ADC. A Cy5 fluorophore (**supplementary Fig.6a**) linker was used because of its biocompatibility and strong fluorescence signal that can be tracked. This enables easy estimation of cellular internalization and localization. The NH_2_-functionalized Cy5-linker was conjugated with the KIM-1 antibody to produce AFC following the similar enzymatic method as mentioned for the ADC (**Fig.4a**). The conjugation reaction of the Cy5-linker was further confirmed by the HIC chromatography (**supplementary Fig.6b**). The peak at 280 nm and 649 nm HIC chromatogram signifies the specific antibody and Cy5 fluorophore conjugation, resulting in exclusively desired homogenous AFC. We confirmed the conjugation specificity in the heavy chain by using SDS-polyacrylamide gel electrophoresis (SDS–PAGE). A denaturing buffer was used to separate the heavy and light chains of the antibody. The presence of the Cy5 fluorescence was observed exclusively at the 50 kDa, corresponding to the selective conjugation in the heavy chain of the antibody (**Fig.4b**). No fluorescence was observed in the light chain of the antibody.

**Figure 4:**
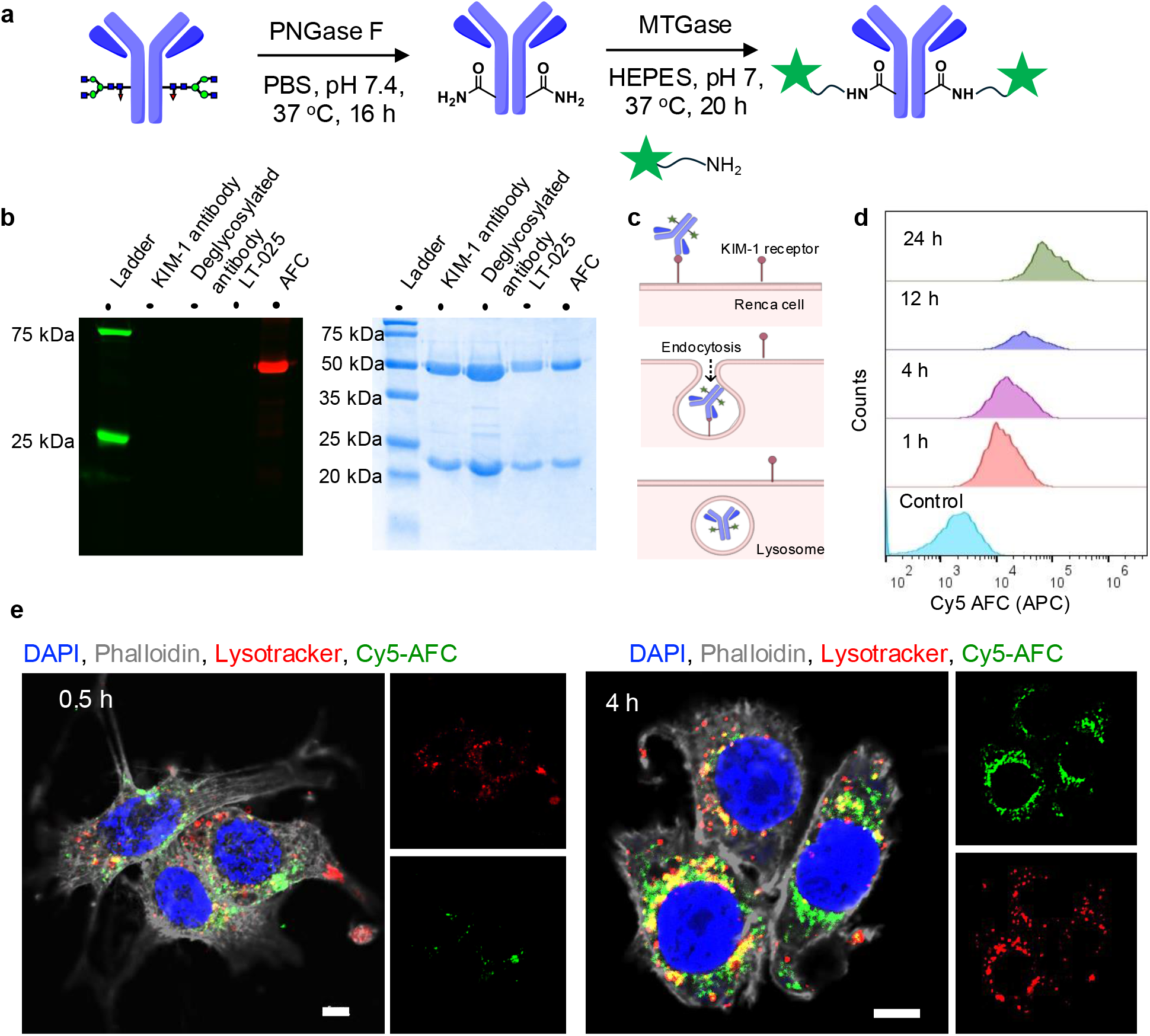
Cellular internalization with KIM-1 engagement. (**a**) Schematic representation of the synthesis of an antibody fluorophore conjugate (AFC) following a similar MTGase reaction as described for the ADC. First, the native monoclonal antibody was deglycosylated in the presence of the PNGase F enzyme, and then the Cy5-fluorophore-linker was attached to the antibody at Q295 in the presence of the MTGase. (**b**) SDS-PAGE (fluorescence imaging and Coomassie staining) of KIM-1 antibody, deglycosylated antibody, LT-025, and Cy5-AFC. The data show attachment of the Cy5 fluorophore in the heavy chain of the antibody. No conjugation of Cy5 was observed in the light chain. (**c**) Schematic representation of the cellular internalization of the ADC and AFC. The AFC first binds with the KIM-1 receptor on the cell surface, and it gets endocytosed into the cell and is transported to the lysosomal compartment. (**d**) Internalization study of Cy5 AFC at different time points. 50 μg/mL of AFC was added to the cell culture. An increase in MFI (Cy5) over time means increased binding/internalization of AFC onto the Renca cell line. (**e**) Confocal microscopy images showing the internalization of the AFC (green) in Renca cells. The images show successful cellular internalization of AFC (50 μg/mL) within 0.5 h and 4 h. An increased cellular internalization of AFC and colocalization at the lysosomal compartment were observed in 4 h. Lysotracker red was used to label the lysosomal compartment. Scale bar = 5 μm.

To test if the ADCs bound to KIM-1 is internalized by endocytosis (**Fig.4c**), we used a metastatic renal cell carcinoma cell line, Renca, which has significant KIM-1 expression (**supplementary Fig.8**). We observed a rapid cellular internalization of AFC in Renca cells using flow cytometry (**Fig.4d**). The histogram represents the increase in the mean fluorescence intensity corresponding to increased cell surface receptor recognition and internalization of AFC with time. However, the flow cytometry data cannot differentiate between the receptor binding on the surface and internalization. Thus, we performed time-dependent confocal microscopy to investigate the cellular internalization and localization. **Fig.4e** shows a considerable amount of cellular internalization of the AFC in Renca cells within 30 minutes and an increased cellular internalization at 4 h. Moreover, we used a lysotracker red dye to evaluate the lysosomal localization of the AFC. We observed localization of the AFC in the lysosomal compartments. Quantification of the colocalization between AFC and lysotracker signal revealed increased colocalization at 4 h compared to 30 minutes (**supplementary Fig.7**), consistent with the hypothesis that antibody payload conjugate first binds with the cell surface receptor, followed by the receptor internalization and localization to the lysosomal compartment.^35^

Next, to confirm that the cellular internalization of the AFC is dependent on the KIM-1 expression on the kidney cancer cell, we generated a KIM-1 overexpressing Renca cell line (Renca^KIM-1++^) by lentiviral transduction. The overexpression of KIM-1 in Renca^KIM-1++^ was characterized using Western blot and flow cytometry analysis. The Renca^KIM-1++^ showed 3 times more KIM-1 expression than wild-type Renca cells (**supplementary Fig.8a**,**b**). Flow cytometric analysis revealed an elevated cellular internalization of AFC in Renca^KIM-1++^ cells compared to Renca cells (**supplementary Fig.8c**). The mean fluorescence intensity (MFI) of AFC showed significantly higher cellular internalization of AFC in the case of Renca^KIM-1++^ than the wild-type Renca, suggesting the antigen-dependent internalization (**supplementary Fig.8d**).

### Cytotoxicity of LT-025, and the synergy with sunitinib, *in vitro*

We next tested the cytotoxicity of LT-025 in Renca cells. As shown in **Fig.5a**, we observed concentration-dependent cytotoxicity in the nanomolar range, with a low *IC*_50_ of 88.67 nM. To investigate the dependency of the cytotoxic efficacy on KIM-1 expression, we next used the Renca^KIM-1++^ cell line. A notably higher cytotoxicity was observed in Renca^KIM-1++^ with a lower *IC*_50_ of 11.78 nM (**Fig.5b**). We evaluated the cytotoxicity of LT-025 on another renal adenocarcinoma cell line RAG, and it showed an IC_50_ of 100.2 nM (**Fig.5c**). Overall, LT-025 showed excellent cytotoxicity with various cell lines, and IC_50_ values suggest the higher potency than many existing ADCs. To confirm that the KIM-1 antibody has minimal binding with non-cancerous cells, we studied the binding effect in T cells and in cocultured cancer cells and T cells. However, a minimal binding was observed in both naïve and activated T cells when compared to the cancer cells (**Fig. 5d,e)**.

**Figure 5:**
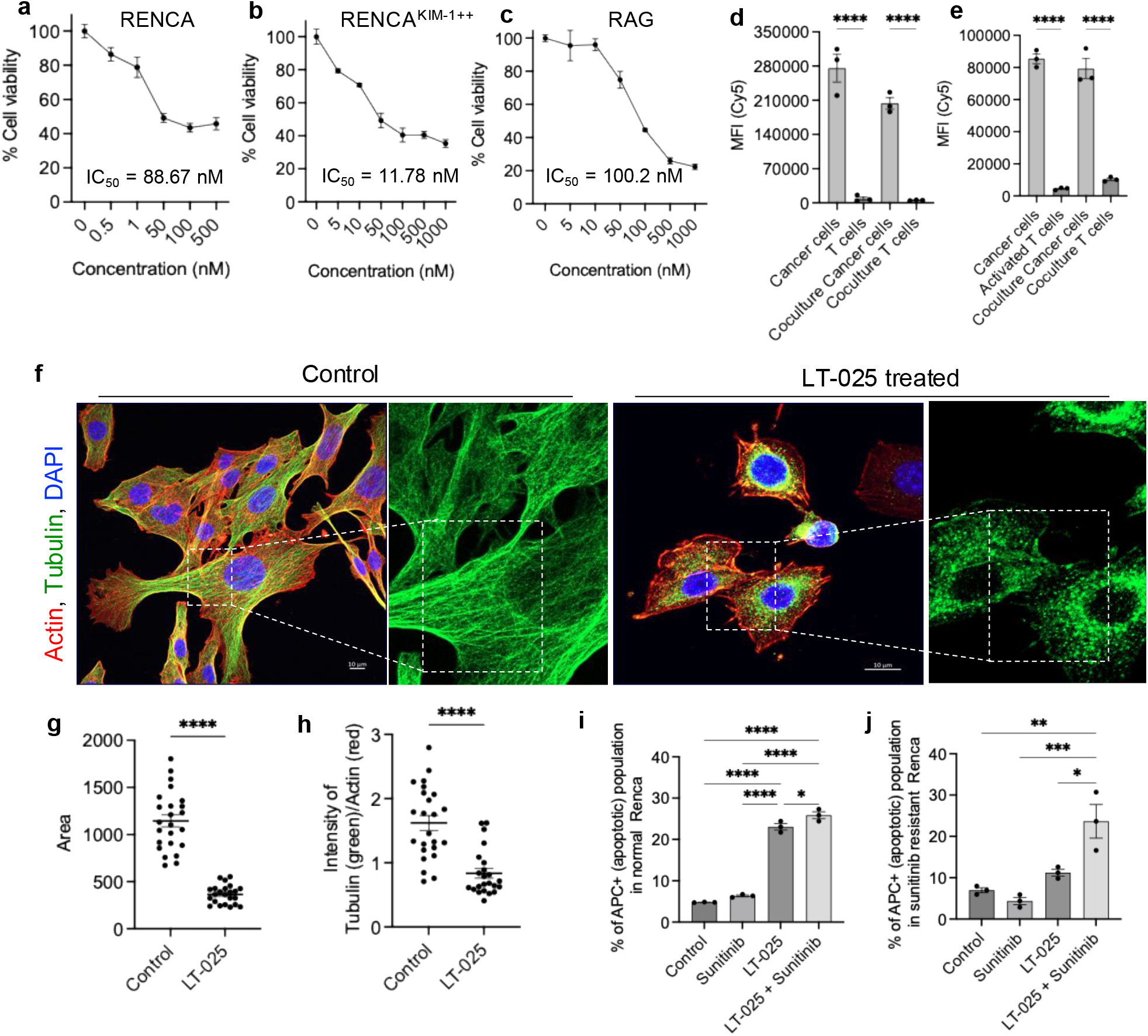
In vitro cytotoxic effect of LT-025. Cell viability and IC_50_ of ADC on (a) normal Renca, (b) Renca^KIM-1++^, and (c) RAG cell lines obtained by MTT assay. The LT-025 exhibits significant cell death with the *IC*_50_ value of 88.67 nM. A relatively lower *IC*_50_ value was observed in Renca^KIM-1++^ th an in normal Renca because of increased receptor-mediated cellular internalization of the ADC. The data were represented as mean ± SEM (n = 3). Mean fluorescence intensity (MFI) of AFC across cancer cells, T cells, and coculture systems. Data compare binding in cancer cells cocultured with (d) naïve T cells and (e) activated T cells. Data are presented as mean ± SEM (n = 3). Significantly higher effect was observed on cancer cells than T cells (f) Confocal images of Renca with and without treatment of LT-025 (100 μg/mL) displaying tubulin disruption. The images showed disruption of the tubulin (green) crosslinking by DM1, a known microtubule inhibitor. Scale bar = 10 μm. (g) Quantification of the area of the control and treated Renca cells. The data showed a significant reduction in the area of the cells. The data were represented as mean ± SEM (n = 23, Unpaired *t*-test with Welch’s correction). (h) Quantification of tubulin in untreated and LT-025 tr eated cells from the confocal images. The data show a reduction in the tubulin intensity upon treatment with LT-025. The mean pixel intensity of the tubulin (green) was plotted after normalizing with the mean pixel intensity of actin (red). The data were represented as mean ± SEM (n = 23, Unpaired *t*-test with Welch’s correction). (i) The bar plot shows the % of apoptotic Renca cells measured with flow cytometry using the Annexin-V AF645 apoptosis kit. The data represent significantly higher apoptosis in normal Renca by LT-025 compared to the control and sunitinib-treated group. 12.5 μM of sunitinib and 100 nM of ADC were used for the study. Data represented as mean ± SEM (n = 3, one-way ANOVA followed by Tukey’s multiple comparison test). The combination of LT-025+sunitinib shows a significantly higher apoptosis of Renca cells. (j) The bar plot showing the % of apoptotic sunitinib-resistant Renca, measured with flow cytometry using Annexin V-AF645 apoptosis kit. The data represent significantly higher apoptosis in sunitinib-resistant Renca by treatment with combination therapy (LT-025+sunitinib) compared to the control and monotherapy. 12.5 μM of sunitinib and 100 nM of ADC were used for the study. Data represented as mean ± SEM (n = 3, one-way ANOVA followed by Tukey’s multiple comparison test).

DM1, the payload used in LT-025, is a known microtubule inhibitor and imposes mitotic arrest followed by cancer cell killing. It prevents microtubule dynamics by binding to the tubulin and converts them to random tubulin aggregates or monomer form^36^. To test whether this mechanism underpins the cytotoxicity of the LT-025, we treated the cancer cells with the ADC and imaged the microtubules using high-resolution confocal microscopy. We observed a substantial reduction of microtubule formation in the Renca cells by ADC (**Fig.5f**). The untreated control Renca cells showed microtubules as a thread-like structure, but the microtubules appeared as an aggregated form in the LT-025 treated cells (**Fig.5f, supplementary Fig.9**), consistent with the mechanism of action of DM1. Moreover, the quantification of such confocal images attributed a significant reduction in the tubulin intensity and the overall cell volume for LT-025 treated cells in comparison to the control (**Fig.5g,h**), consistent with cell death.

Targeted therapeutics, such as sunitinib, have been a mainstay in the treatment of metastatic RCC. Indeed, the combination of targeted therapeutics and immune checkpoint inhibition has emerged as the first line is the current treatment for metastatic RCC (mRCC) patients, and sunitinib-monotherapy is still the option for patients who are ineligible or unresponsive to ICI. However, most patients experience disease progression after initial response to treatment and tumor shrinkage. Given the non-overlapping mechanism of action, we rationalized LT-025 could augment sunitinib efficacy. Indeed, as shown in **Fig.5i**, significantly higher induction of apoptosis was seen in the LT-025+sunitinib treated cells compared to the untreated control and monotherapy. Next, we have generated a sunitinib-resistant Renca cell line by increasing the concentration of sunitinib according to the reported procedure.^37^ We investigated the effect of LT-025 on sunitinib-resistant cells, alone and in combination with sunitinib. The combination of LT-025+sunitinib showed a significantly enhanced apoptosis induction compared to monotherapies in the sunitinib-resistant Renca cells (**Fig.5j**). Hence, the combination of LT-025+sunitinib has the potential to offer enhanced therapeutic outcomes in RCC.

### LT-025 synergizes with sunitinib *in vivo*

Prior heterogenous KIM-1 targeted ADCs were limited by toxicity.^14^ As a first step, we performed a detailed toxicity evaluation using repeat dose escalation in immunocompetent balb/c mice. We injected LT-025 (every 3-4 day) at increasing doses of 0.83 to 3.33 mg/kg. We sacrificed 5 mice after the 1.67 mg/kg dose and the rest after the 3.33 mg/kg dose to evaluate for any toxicity. No significant body weight loss or clinical signs of toxicity were observed at any of the dose levels (**Fig.6a**). We performed detailed biochemistry analysis for liver and kidney functions. No abnormal change in blood alkaline phosphatase (ALP), aspartate aminotransferase (AST), alanine aminotransferase (ALT), Creatine kinase, total protein, globulin, albumin, bilirubin, and blood urea nitrogen (BUN) was observed at either of the doses used (**Fig.6b, supplementary Fig.10**). For the assessment of potential nephrotoxicity, we performed hematoxylin and eosin (H&E) staining and periodic acid-Schiff (PAS() staining on kidney tissue samples collected after treatment with LT-025 at different doses. The preserved glomerular structure and distribution, along with intact tubular morphology and normal renal architecture, indicate the absence of significant histopathological renal toxicity in the tested doses (**Supplementary Fig.11, 12**).

Next, we used the syngeneic Renca tumor model in balb/c mice to evaluate the tumor growth inhibition by LT-025 as a monotherapy and in combination with sunitinib. A subcutaneous tumor was implanted in the right flank of the male balb/c mice. The mice were monitored for their body weight and tumor growth. We randomized the mice into 4 groups once the tumor reached 50-75 mm^3^ in volume. Mice were dosed with 1.67 mg/kg of LT-025 on days 8, 12, 17, and 21 post tumor implantation, using intravenous injection. Sunitinib (20 mg/kg) was given orally on days 7, 11, 16, and 20 (**Fig.6c**). While both sunitinib- and LT-025-treated groups showed reduced tumor growth than control mice (**Fig.6d**), the combination of LT-025 and sunitinib resulted in a synergistic increase in therapeutic efficacy compared to monotherapies. The combination did not result in any significant treatment-related toxicity.

**Figure 6:**
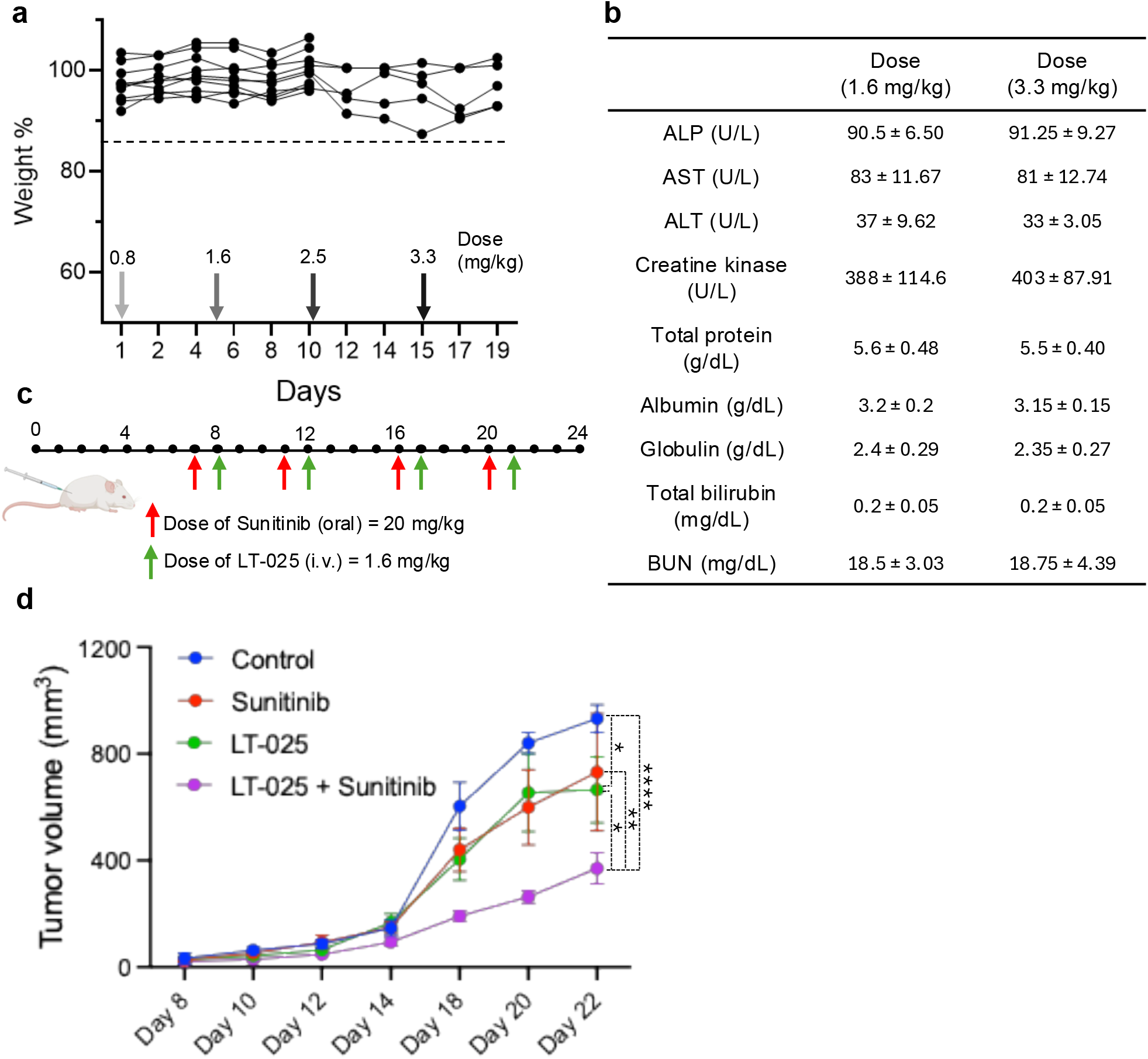
LT-025 synergizes therapeutic efficacy with sunitinib. (**a**) Dose escalation study to evaluate any treatment-related toxicity in mice by LT-025. The data shows no significant change in the body weight of each mouse during the treatment. We sacrificed 5 mice after the 1.6 mg/kg dose and the remaining after 3.3 mg/kg to perform blood biochemistry analysis. No significant toxicity, i.e, no change of body weight more than 20% of the initial weight has been observed. (**b**) Blood biochemistry analysis from the dose escalation study after the doses of 1.6 and 3.3 mg/kg. The data show no abnormal levels of critical liver and kidney enzymes and blood components, signifying safe treatment of the LT-025. (**c**) The schematic shows the study design for evaluating the therapeutic efficacy of LT-025 using a subcutaneous Renca tumor model in syngenic balb/c mice. The injection regime is indicated using red (sunitinib) and green (LT-025) arrows. We followed the standard mode of administration of ADC and sunitinib. (**d**) Tumor growth curve showing the change in tumor volume of each treatment group. The data show a significant reduction in the tumor volume by LT-025. The therapeutic efficacy of LT-025 increases significantly in the presence of sunitinib compared to monotherapies. The data were represented as mean ± SEM (n = 5, two-way ANOVA followed by Tukey’s multiple comparison test).

## DISCUSSION

Mortality in advanced RCC patients remains high because of the lack of effective treatment approaches. While antibody drug conjugates have emerged as a leading class of therapeutic agents for treating cancer, there are no ADCs approved for RCC. In this study, we described the engineering of a novel KIM-1 targeting antibody drug conjugate for use in RCC. Our study shows the combination of a KIM-1 targeting antibody drug conjugate, LT-025, could improve the therapeutic outcome in combination with sunitinib.

Patient biopsy analysis revealed that the majority (70%) of the patients have significantly higher KIM-1 expression in cancer tissue than in the adjacent normal tissue, consistent with published reports^20–23^, making KIM-1 a promising target for the ADC design for kidney cancer. Unfortunately, prior efforts at developing KIM-1 targeting ADCs failed due to heterogeneity of the product and premature release of the payload from the antibody and resulting off-targeting toxicity.^14^ We used an enzymatic transamidation reaction in specific glutamine residues in the Fc region of the native antibody. This resulted in a precise DAR of 2 in LT-025, i.e, one drug conjugation per heavy chain. This conjugation at the heavy chain is also verified in the case of AFC. The HIC chromatogram showed a single major product, signifying that the reaction yields homogenous product. This eliminates the requirement of calculating the average DAR from multiple peaks of the HIC chromatogram, a common method followed in the case of heterogeneous ADC.^38^ The conjugation followed a two-step process: (i) deglycosylation of the antibody using PNGaseF enzyme, (ii) MTGase-based conjugation of the NH_2_-functionalized drug-linker with Q295 of the heavy chain of the anti-KIM-1 IgG1 subtype antibody. The regioselectivity of the conjugation on the Q295 over other glutamine residues is likely governed by (i) the availability of the glutamine on the surface for the enzyme to bind and (ii) the flexibility of the region known as the B-factor.^39^ However, in the native IgG1 antibody, the Q295 position is sterically hindered because of the presence of the glycan at N297. PNGase-F based method of deglycosylation selectively removes the polysaccharide unit from the N297 position.^30,31^ The removal of the Fc glycan not only offers increased accessibility of the Q295, also increases the flexibility of the region.^40,41^ Fontana et al. have shown the relation of the flexibility of the polypeptide chain with the selectivity of the MTGase reaction.^39^

Apart from site-specific conjugation, the use of a non-cleavable linker, as is the case in this study, to conjugate the drug to the antibody can lower premature release, reducing the off-target toxicity.^42^ For example, Trastuzumab DM1 (T-DM1, Kadcyla), a clinically approved ADC, is constructed with a non-cleavable linker and has shown successful payload delivery to the tumor cells.^43^ However, such a construct needs to be internalized, where the lysosomal environment can break down the ADC and release the potent payload. Indeed, we observed that LT-025 showed identical antigen binding as the KIM-1 antibody, and was rapidly internalized into the cell dependent on KIM-1 expression, and exhibited potency in the nanomolar concentration range *in vitro*. Taken together, the increased stability and the homogeneity could explain the improved safety of the ADC *in vivo*. Additionally, in preclinical efficacy studies, monotherapy with LT-025 was as effective as sunitinib, which is the standard of care in the clinic for advanced RCC that have progressed on combination therapies.^44,45^ Interestingly, the combination of sunitinib and LT-025 resulted in a synergistic anti-tumor outcome than either agent given as monotherapy. It should be noted that monotherapy is rarely effective in RCC, and the standard of care is combination therapies. Unfortunately, the combinations of sunitinib with other kinase inhibitors did not show a promising outcome in RCC.^46,47^ Hence, our results attain significance as it indicates that KIM-1 DM1 ADC could potentially be added to sunitinib in the clinics without additional toxicity.

Further studies are required to translate to humans, including using a humanized anti KIM1 antibody and safety studies in additional animal models. Additionally, the ADC can be modified to deliver both DM1 and sunitinib, which can further optimize the therapeutic index. In conclusion, our study indicates that a homogenous and stable KIM-1 targeting ADC can offer a promising therapeutic option for RCC, alone or in combination with targeted therapeutics.

## MATERIALS AND METHODS

### Synthesis of the drug-linker

In a clean and dry round-bottom flask, Mal-amido-PEG_9_-NH_2_ (Broadpharm, 0.020 mmol) was mixed with acetonitrile under an inert atmosphere. N, N-Diisopropylethylamine (DIPEA, 0.1 mmol) was added, followed by the addition of DM1 (MedChemExpress, 0.024 mmol). The mixture was stirred at room temperature for 3 hours. After the reaction, acetonitrile was evaporated in vacuo, and the residue was extracted using ethyl acetate and water. The organic layer was collected and dried over Na_2_SO_4_. The residue was washed with 20% ethyl acetate and hexane to remove the excess unreacted DM1. The expected compound was collected, dried under vacuum, and characterized by NMR and mass spectrometry. The product was characterized by ^1^H NMR and ^13^C NMR using Bruker Avance-III HD Nanobay spectrometer, 400.09 MHz. ^1^H-NMR spectrum (400 MHz, CDCl_3_) of DM1 linker. δ (ppm) 0.81 (s, 3H), 1.437 (s, 3H), 1.47 (s, 3H), 1.57 (t, 2H), 1.60 (t, 2H), 1.66 (s, 3H), 2.11 (s, 3H), 2.22 (d, 1H), 2.59 (q, 1H), 2.64 (d, 2H), 2.87 (s, 3H), 2.9 (t, 2H), 3.04 (d, 3H), 3.10 (t, 2H), 3.22 (s, 3H), 3.38 (s, 3H), 3.43 (t, 2H), 3.52 (t, 1H) 3.69 (m, 32H), 3.81 (q, 2H), 3.92 (t, 1H), 4.01 (s, 3H), 4.30 (t, 1H), 4.79 (d, 1H), 5.39 (q, 1H), 5.61 (t, 1H), 6.3 (d, 1H), 6.45 (t, 1H), 6.65 (s, 1H), 6.84 (s, 1H). ^13^C NMR (101 MHz, CDCl_3_) of DM1 linker δ (ppm) 176.64, 174.54, 170.82, 170.45, 168.86, 161.84, 156.07, 152.35, 142.23, 141.21, 139.34, 133.37, 127.65, 125.34, 122.23, 118.82, 113.27, 88.65, 80.97, 78.31, 74.23, 70.57, 70.01, 69.84, 60.06, 56.69, 53.97, 52.47, 46.73, 42.05, 40.25, 39.94, 38.99, 37.81, 35.95, 35.60, 34.31, 33.46, 32.54, 30.82, 26.88, 19.91, 18.70, 17.58, 15.64, 14.70, 13.50, 12.30, 11.92. The liquid chromatography and mass spectrometry (LC-MS) of the product was carried out in Agilent 6125B mass spectrometer attached to an Agilent 1260 Infinity LC. Calculated mass for [M+H]^+^: 1345.6 Da, observed mass: 1345.6 Da. A gradient method of 100% water to 100% acetonitrile in the presence of +0.1% formic acid was used for HPLC using a C18 column.

### Synthesis of ADC (LT-025)

#### Deglycosylation

KIM-1 antibody (anti-mouse, Rat IgG1, BioXcell, 2 mg/mL in PBS) was taken and added 2 μL of PNGase F (New England Biolabs, 1000 units). Incubated at 37 °C for 20 h. The solution was filtered through 50 kDa Amicon filters three times using PBS and diluted to a 10 mg/mL concentration.

#### Antibody-Linker Conjugation

Deglycosylated KIM-1 antibody (5 mg/mL in 1:2 PBS and HEPES buffer) was incubated with the linker (18.5 μl, 50 eq, 20 mg/mL in DMSO) and enhanced transglutaminase enzyme (Sigma Aldrich, 22.7 μl, 2.5 eq) at 37 °C for 20 h. The antibody-drug conjugate formed was purified using 50 kDa Amicon filters by washing with PBS five times. The product was purified from the trace amount of remaining enzyme using protein A agarose beads (Santacruz, 200 μL). After the conjugation reaction, protein A agarose solution was added and incubated at 4 °C for 1 h. The beads were washed with PBS three times in a mini spin column (QIAmp), and then added 0.1 M glycine, pH: 3.2 solution to elute the ADC. This was further subjected to buffer exchange with PBS, pH 7.4, using Amicon filters, and the product LT-025 was stored at 4 °C. The final concentration of the ADC was measured using a Thermoscientific NanoDrop One.

### Synthesis of Cy5-AFC

The deglycosylated KIM-1 antibody (5 mg/mL) was diluted in 1:2 PBS and HEPES buffer and mixed with Cy5 linker [N-(m-PEG4)-N’(amino-PEG3)-Cy5, Broadpharm, BP-23008, 40 eq., 8.7 μL from 20 mg/mL stock solution in water] and enhanced transglutaminase enzyme (22.7 μL). The mixture was incubated at 37 °C for 20 h in the dark. The antibody-fluorophore conjugate (AFC) formed was purified using 50 kDa Amicon filters by washing with PBS five times.

### Characterization of ADC using Hydrophobic Interaction Column Chromatography

The conjugation efficiency was monitored using hydrophobic interaction column (HIC) chromatography using Agilent 1200 HPLC system connected with a Butyl-NPR column (3.5 cm × 4.6 mm, 2.5 μm particle size). Mobile phase A: 25 mM phosphate buffer pH 7 containing 1.5 M ammonium sulfate, and B: 25 mM phosphate buffer pH 7 containing 20% isopropanol, were run at a gradient from 0% to 100% B at the flow rate of 0.5 mL/min.

### Electron Spray Ionization-Mass Spectrometry (ESI-MS)

The antibody and ADC samples were reduced with dithiothreitol (DTT) to separate into heavy and light chains. The solution was then washed and desalted using 3000 MWCO Millipore Amicon filters. 5 mg of sample was directly injected (bypassing column) into the Agilent 1100 High-Performance Liquid Chromatography (HPLC) connected with a Bruker Maxis Impact LC-q-TOF Mass Spectrometer in the presence of the solvents acetonitrile and water with 0.1% formic acid. We have recorded a mass range from 300 Da to 3000 Da. And the signals were deconvoluted to obtain the total mass of the reduced antibody.

### Immunohistochemistry and image analysis

Tissue microarrays (TMA) for RCC patients’ biopsies (KD321a) were obtained from TissueArray.com. The TMA has cancer and matched adjacent normal kidney tissue from patients randomized based on age, sex, and stage of the cancer. The FFPE tissue sections were deparaffinized by dipping into xylene and rehydrated by the gradient of the ethanol-water mixture. The antigen retrieval of the tissue specimen was performed using citrate and Tris buffer at 95 °C. The blocking step was done by 5% bovine serum albumin, and 2% goat serum. Antibody staining was performed by specific primary antibody anti-human-KIM-1 (Cell Signalling Technology, Cat. No. 14971, 1:100 dilution), followed by secondary antibodies (anti-rabbit AF488, Thermo Fisher Scientific, Cat. No. A11008, 1:1000 dilution). Images were acquired in a TissueFAXS Plus slide scanner (TissueGnostics USA) equipped with a Hamamatsu Orca Flash 4.0 V2 cooled digital CMOS camera and Zeiss 20x EC Plan-NEOFLUAR 0.5NA air objective. Each image of the tissue specimens is composed of 16 individual image view fields. The quantification of KIM-1 was carried out by measuring the mean pixel intensity using Image*J* software and normalizing with the mean pixel intensity of DAPI.

### Cell culture

Murine renal cell carcinoma, Renca (ATCC, CRL-2947), was cultured in RPMI-1640 (ATCC), and renal adenocarcinoma cell RAG (ATCC, CCL-142) was cultured in EMEM (ATCC) supplemented with 10% v/v heat-inactivated fetal bovine serum (FBS) (Thermo Fisher Scientific) and 1% v/v Penicillin-Streptomycin (Thermo Fisher Scientific). Cells were cultured, maintained in sterile conditions, and housed in a water-jacketed CO_2_ incubator with an atmosphere at 37 °C and 5% CO_2_. Cells were routinely tested and mycoplasma-free.

### Flow cytometry for cellular internalization

Renca cells were seeded in a culture-treated 24-well plate (100k cells per well) and incubated at 37 °C under 5% CO_2_ for 24 h. The AFC sample (50 μg/mL) was added to the fresh media after a wash with DPBS and incubated under the same conditions for different time points, such as 1 h, 4 h, 12 h, and 24 h. After the time, the cells were washed with DPBS two times and collected after trypsinization. The collected cells were resuspended in 300 μL of cell staining buffer to run the flow cytometry. An Agilent Novocyte flow cytometer was used, and the data were analyzed using FlowJo.

### Immunostaining, fluorescence, and confocal microscopy

Renca cells were seeded on sterile 12 mm coverslips in a 24-well plate (100k cells per well) and incubated at 37 °C under 5% CO_2_ for 16 h. The AFC sample (50 μg/mL) was added to the fresh media after a wash with DPBS and incubated under the same conditions for different time-points, such as 0.5 h, 4 h. The cells were washed, and 500 nM of Lysotracker dye (red, Invitrogen) was added and incubated for 30 min. Later, the cells were fixed with 4% paraformaldehyde (Electron Microscopy Sciences), followed by staining with Phalloidin 488 (green, Invitrogen, 1: 400) at an incubation time of 1 h. The nucleus staining was carried out using Hoechst (blue, Invitrogen, 1:200). The coverslips were then mounted on the glass slide with Prolong Glass Antifade Mounting Media.

To study the effect of LT-025, a 100 µg/mL solution of ADC in fresh complete medium was added to adhered cells on the coverslips and incubated for 24 h. Untreated cells were also taken in similar conditions as the control. After washing, the cells were fixed in 4% paraformaldehyde (Electron Microscopy Sciences), followed by permeabilization and blocking in PBS containing 5% goat serum and 0.3% Triton X for 1 h at room temperature. The primary antibody anti-α,β tubulin antibody (cell signaling technology, rabbit IgG, 1:50) was incubated for 16 h at 4 °C. After washing, Alexa Fluor 488 secondary antibody (green, Invitrogen, anti-rabbit IgG, 1:500) in 1 % BSA and 0.3 % Triton X was added and incubated for 1 h. Further F-actin staining was carried out using Rhodamine-Phalloidin (red, Invitrogen, 1:400) for 1 h at room temperature. Finally, the nucleus staining was carried out using Hoechst (blue, Invitrogen, 1:200). The coverslips were then mounted on the glass slide with Prolong Glass Antifade Mounting Media. Imaging was done using either Zeiss LSM 800, Airyscan confocal laser scanner microscope with Zen 2.3 software. Postprocessing of the image was done in ImageJ v1.52a software.

### Enzyme-linked immunosorbent assay (ELISA)

The antigen-binding ability of ADC was studied by Enzyme-Linked Immuno-Sorbent Assay (ELISA). 50 μl solution 2 μg/mL solution (100 ng) of mouse KIM-1 (SinoBiological, 50321-M16H) in PBS was coated on an ELISA plate (ELISA Max, Biolegend) and incubated overnight at 4 °C. The plate was washed using PBST three times for five minutes each. Then, 300 μl of 10% BSA was added for blocking and incubated at room temperature for 90 min. Later, 100 μl of LT-025 and KIM-1 antibody control was added in varied concentrations and incubated at 37 °C for 1 h. After subsequent washing, HRP conjugated secondary antibody of IgG1 (1:1500 dilution) was added and incubated for 30 minutes at 37 °C. TMB substrate was then added after multiple washes with PBST. After five minutes, 2 N H_2_SO_4_ was added to stop the reaction, and the absorbance was recorded at 450 nm using Epoch Biotek microplate reader. The data was analyzed using GraphPad Prism. The dissociation constant K_d_ was calculated from the absorbance versus concentration plot fitted to a one-site specific binding equation using GraphPad Prism software.

### Biolayer Interferometry

Octet RH16 (previously Octet 698 RED384) from Sartorius was used for the BLI measurement. The analysis was carried out in a 699 Octet Red 384-well plate. PBS with 0.05% Tween 20 (PBST) was used for the hydration of the streptavidin sensor. The sensors were hydrated for two minutes before analysis. PBST was also used as the analysis buffer and for sample preparation to minimize all possible non-specific interactions. Mouse-KIM-1 antigen (SinoBiological, 50321-M16H) was biotinylated using Thermo Scientific EZ-Link™ Micro Sulfo-NHS-LC-Biotinylation Kit. 10 μg/mL of biotinylated KIM-1 was used as the load. After the baseline setting for 120 seconds, loading was performed for 120 seconds, followed by another baseline setting for 120 seconds. The association and dissociation with the LT-025 samples were recorded for 600 seconds each. Negative control for sensors was carried out without loading the biotinylated protein, and a null concentration of samples was used as a negative control for the samples, and these responses were subtracted from the samples. The analysis of the result was carried out in Octet Analysis Studio 13.0. Inter-step correction was performed by aligning to the baseline step, and noise filtering was performed.

### Stability of ADC

The stability of the ADC was determined using the HIC chromatography method. 1 mg/mL of LT-025 in mouse plasma was incubated at 4 °C and room temperature. 10 μL of each sample was injected into the HPLC connected to the HIC. The area under the curve of the peak corresponding to the ADC was measured and plotted for different days.

### DAR calculation

The DAR of LT-025 was calculated using UV-visible spectroscopy. First, the extinction coefficient of both the antibody and the linker was calculated using Beer-Lambert’s law with a known concentration. Then the concentration of antibody part (C_mAb_) and drug-linker part (C_drug_) of the ADC was calculated using equations 1, and 2, and DAR was calculated using equation 3.

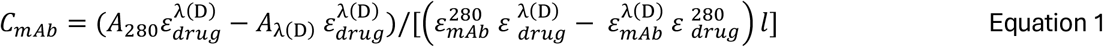

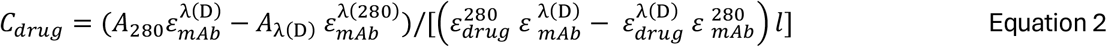

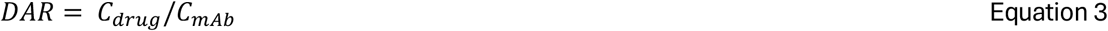

where, *A*_280_ is the absorbance of ADC at 280 nm, 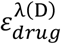 is the extinction coefficient of drug-linker at 254 nm, *A*_*λ*(D)_ is the absorbance of ADC at 254 nm, 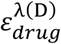 is the extinction coefficient of drug-linker 254 nm, 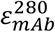 is the extinction coefficient of antibody at 280 nm, 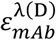 is the extinction coefficient of antibody at 254 nm, 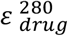 is the extinction coefficient of drug-linker at 280 nm and *l* is the pathlength, which is 1 cm.

### In-vitro cytotoxicity studies

Renca/RAG cells were seeded in a culture-treated 96-well plate (2.5 k cells per well) and incubated at 37 °C under 5% CO_2_ for 24 h. The LT-025 sample was added in increasing concentration in the fresh media and incubated under the same conditions for 48 h. Further, the ADC-containing media were removed from the wells, and 0.5 mg/mL MTT dissolved in basal medium (100 μL) was added and incubated for 3 h. Later, the formazan crystals were dissolved in DMSO by replacing the MTT solution. The absorbance at 570 nm was read using a BioTek Epoch microplate reader. The *IC*_50_ values were then calculated by fitting the data with the logistic function to create a sigmoidal dose-response curve. All measurements were performed in triplicate. Data were analyzed by Prism 10 (GraphPad) software.

### Generation of Renca^KIM-1++^ by overexpressing KIM-1

We have used lentiviral transduction of wild-type Renca cells to overexpress KIM-1. HAVCR-1 Mouse tagged ORF Clone Lentiviral particle (MR203831L3V, Origene, Rockville, MD) containing the desired murine-KIM-1/HAVCR1 gene along with a puromycin-resistant gene for selection, was used to transfect mouse renal adenocarcinoma cells (Renca). The transduction of lentivirus was performed according to the manufacturer’s protocol. A similar lentivirus with the same ORF but lacking the KIM-1 gene (PS100092V) was used as a negative control. Antibiotic selection using 0.5 μg/mL puromycin in complete RPMI medium was performed (according to the manufacturer’s recommendation) to enrich the Renca^KIM-1++^ population. The concentration of puromycin was determined by generating a kill-curve using MTT assay with the transduced Renca cells. The KIM-1 expression in Renca^KIM-1++^ was investigated using western blot and flow cytometric analysis.

### Western blot

Normal Renca and Renca^KIM-1++^ cells were seeded in a 6-well plate at a density of 600k cells per well and incubated at 37 °C under 5% xCO_2_ for 16 h. The Cells were lysed with RIPA lysis buffer supplemented with protease and phosphatase inhibitors for 30 min at 4 °C with mild vortexing every 10-15 min, followed by centrifugation. The amount of protein was optimized by bicinchoninic acid assay (Thermo Scientific) according to the manufacturer’s protocol. 40 µg of protein lysates were electrophoresed on a 10% polyacrylamide gel at 80-120 V until resolved completely. The proteins were transferred to an Immun-Blot PVDF membrane (Biorad, 0.2 µm) through wet blotting for 3 h at 300 mA. Membranes were blocked by either 5% non-fat dry milk in TBS-T (TBS containing 0.1% Tween-20) or 5% BSA in 1x TBS-T for 1h and incubated with appropriate primary antibodies at 4 °C for 16-18 h. After that, the membrane was washed 3 times with 1x TBS-T. Then the membrane was incubated with secondary antibody HRP-goat-anti-mouse/rabbit (1:1000) (BioRad) for 1 h at room temperature. The membranes were washed two times with 1x TBS-T and one time with TBS before being developed using femto chemiluminescent substrate (Thermo Scientific) and imaged using Biorad ChemiDOC XRS+ using Image Lab version 6.1 image analysis. The antibodies used for the western blot analysis were anti-mouse-KIM-1 antibody (Abcam, rabbit polyclonal, 1:500), E-Cadherin (Cell Signaling Technology, Cat. No. 3195, 1:1000), or GAPDH (Cell Signaling Technology, 1:1000)

For the study of expression of E-cadherin, different concentrations of LT-025 from 0 nM to 500 nM were treated to 600k cells adhered on 6-well plates in fresh complete media for 24 h. After the treatment, the media was removed, and the cell lysate was prepared. The western blotting was carried out according to the same protocol mentioned above.

### Development of sunitinib-resistant cells and apoptosis assay

For the development of sunitinib-resistant Renca cells, the parental cells were continuously treated with sunitinib in increasing concentrations of sunitinib from 1 μM, 2.5 μM, 5 μM, 6 μM, 7.5 μM, 8.5 μM, 10 μM to 12.5 μM. The cells were allowed to grow in sunitinib-containing media until they were confluent. Then the cells were split into two flask: one with the same concentration of sunitinib where the cells were already in, and another with the higher dose. The cells followed a pattern of getting confluency in 4 days, and the treatment continued for one month. After the dose of 12.5 μM of sunitinib, the growth of cells was reduced considerably, which is a characteristic of resistant cells.

For the apoptotic assay, 100 k of these sunitinib-resistant cells and normal Renca cells were treated with 100 nM of LT-025, 12.5 μM of sunitinib, and a combination of both for 24 hours in a 24-well plate. The cells were collected, washed, and stained with Annexin V-AF645 (Invitrogen, A23204) for 15 minutes at RT. The cells were then diluted with 200 μl of binding buffer and run on the flow cytometry with Novocyte flow cytometry. The analysis was carried out using FlowJo 10.10.0, and the data were plotted in GraphPad Prism.

### In vivo dose escalation study

We have performed a dose escalation study in 10 male balb/c mice (8-10 weeks old, Charles River Laboratories, Strain Code 028). The LT-025 was administered by i.v. injection at the dose of 0.83 mg/kg to 3.33 mg/kg at an interval of 3-4 days. The mice were monitored for their weight and normal movement to evaluate any signs of toxicity. We have sacrificed 5 animals after the 1.67 mg/kg dose and the remaining 5 after the 3.33 mg/kg dose. We have collected blood from each mouse by intracardiac puncture and performed detailed blood biochemistry analysis for critical components representative of any systemic toxicity.

### In vivo tumor reduction studies

Renca cells (1×10^6^ cells in PBS) were injected subcutaneously into flank region of 8-10 weeks syngeneic male balb/c mice (Charles River Laboratories, Strain Code 028). The animals were randomized into four different treatment groups once the tumor size is in between ~50 - 100 mm^3^. Each group has 5 animals. LT-025 was administered at the dose of 1.67 mg/kg via the tail vein. Sunitinib (20 mg/kg) was given orally in a formulation with N-carboxymethyl cellulose and Tween 80. The drug administration was followed according to the schedule in Fig.6c. The mice’s weight and tumor volume were monitored every other day.

The tumor volume was calculated using **Equation 4**:

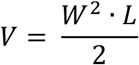

where *V, W*, and *L* are the tumor volume and the shortest and the longest diameter, respectively, as measured using a vernier caliper. The animals were euthanized when it almost reached the permissive tumor volume. The blood samples were collected for serum toxicity analysis on the day of euthanasia. All procedures were performed according to the approved animal protocol by the BWH Institutional Use and Care of Animals Committee.

### Statistical analysis

All statistical analysis was performed using Prism 10 (GraphPad) software. Experimental data were expressed as means ± SEM (otherwise specifically stated), and a *t*-test or one- or two-way analysis of variance (ANOVA) followed by an appropriate post-test was used to calculate statistical significance. *P* < 0.05 was considered significant and represented as *. Other P values were represented as \*\**P* < 0.01, \*\*\**P* < 0.001, and \*\*\*\**P* < 0.0001.

## ACKNOWLEDGMENTS

We acknowledge the Harvard Center for Mass Spectrometry for the guidance on data acquisition and analysis. We also acknowledge Sreyan Ghosh and Natália Vadovičová for helping with the preliminary studies.

## Author contributions

Conceptualization: T.S., Methodology: L.T., K.V., B.S., and A.K. Investigation: L.T., K.V., B.S., A.K., and T.S. Supervision: T.S. and S.S. Writing—original draft: T.S., S.S., and L.T. Writing—review and editing: T.S., L.T., B.S., and S.S.

