## Supplemental Data for "Combining a homogenous KIM-1-DM1 antibody drug conjugate with sunitinib in renal cell carcinoma"

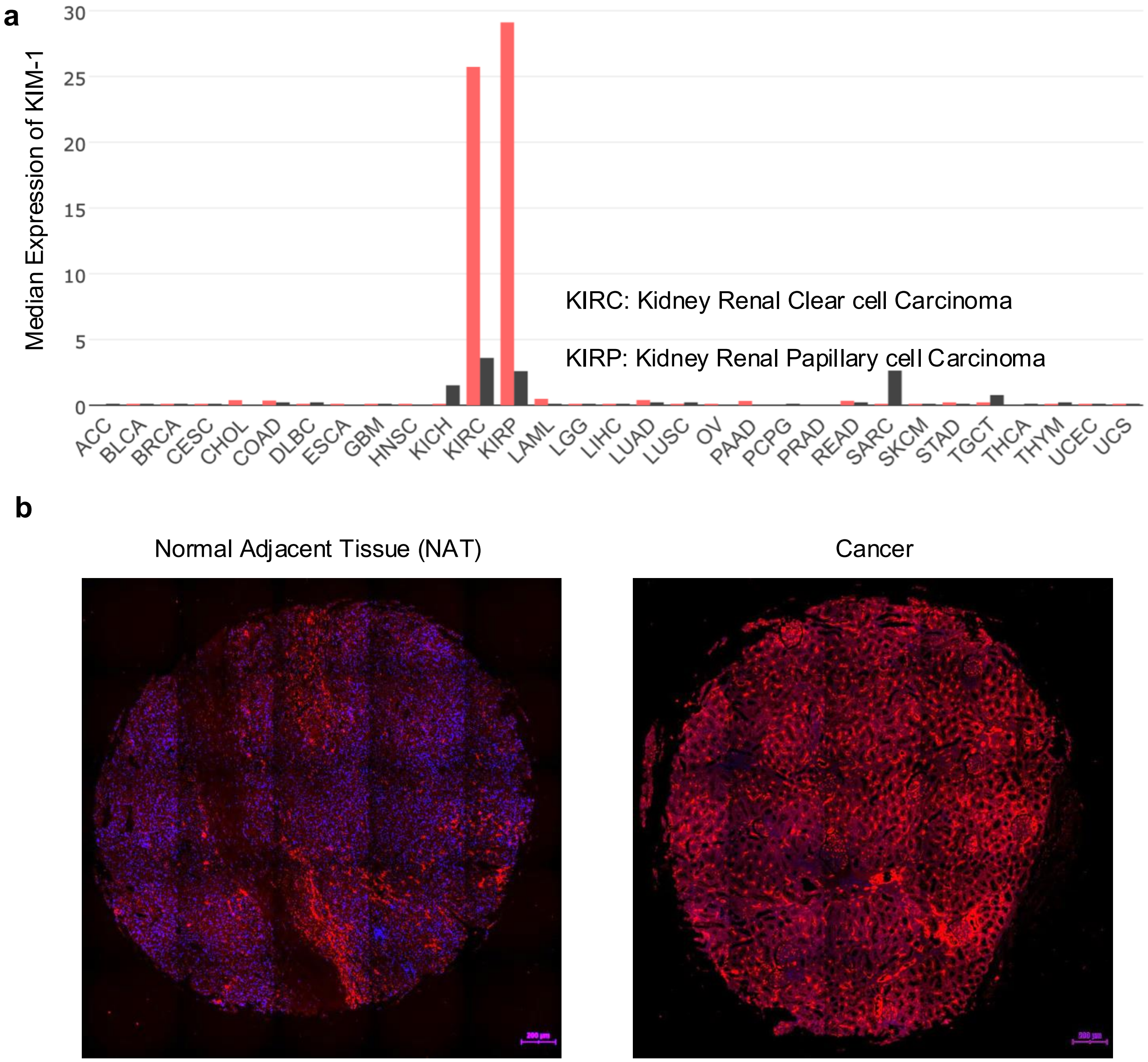


**Supplementary Figure 1:** (**a**) Median expression of the KIM-1 gene in different cancer and normal tissues obtained from the TCGA dataset using Gene Expression Profiling Interactive Analysis (GEPIA). (**b**) Representative images for immunostaining of kidney tissue microarray by anti-KIM-1 (red) antibody and DAPI (blue). Tissue specimens were collected from randomized patients based on age, sex, and stage of cancer. The normal adjacent tissue (NAT) and tumor tissue were collected from the same patients. The images presented without field correction show that a total of 25 images were combined to form the entire circular image.

a

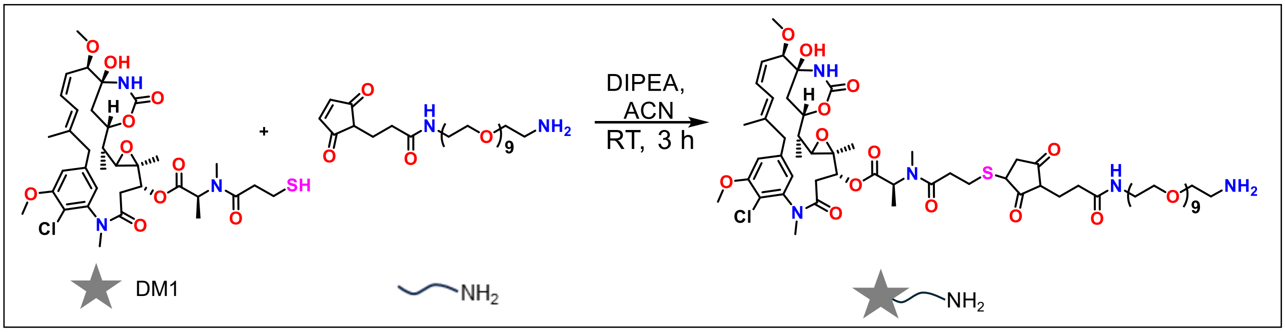


b

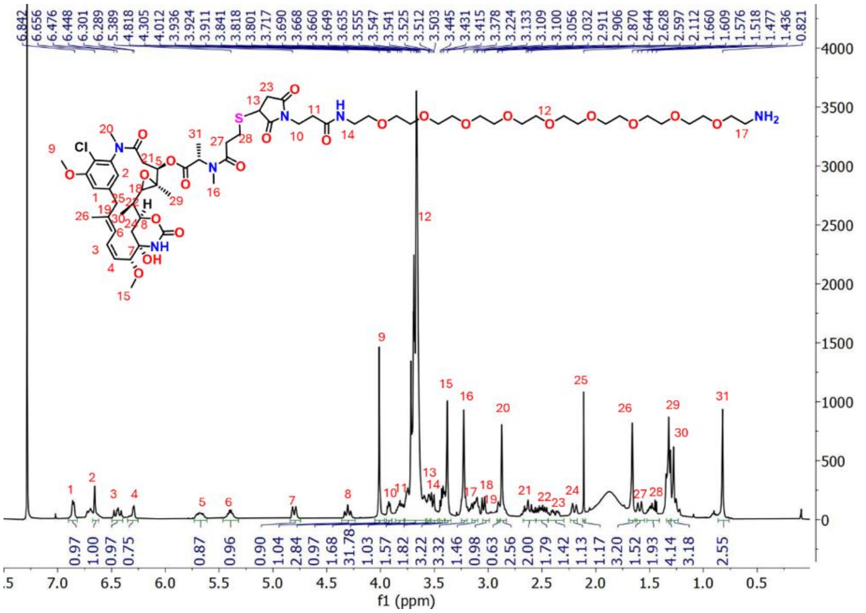


c

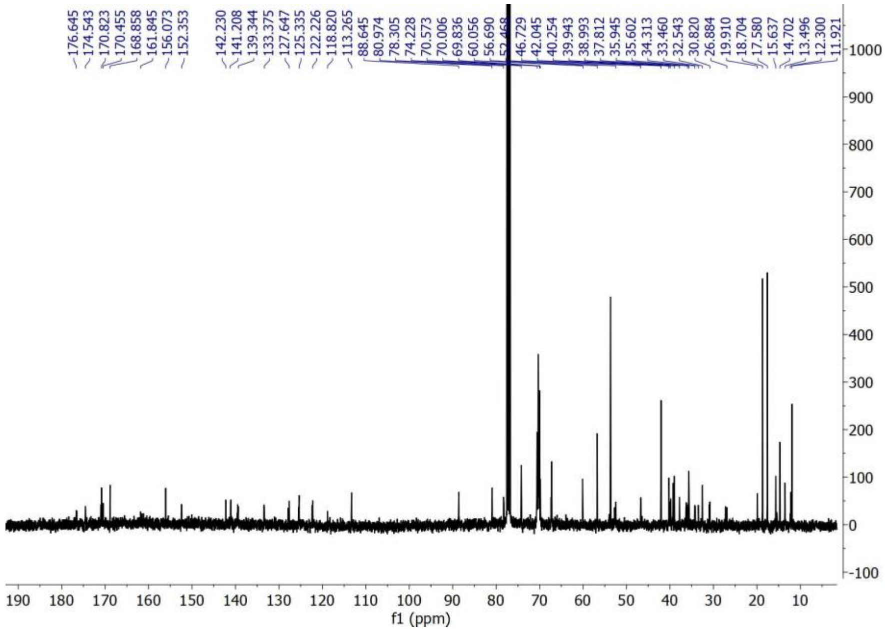


**Supplementary Figure 2:** (**a**) Schematic representation of the DM1-linker preparation in a single step using DM1 and malamido-PEG$_9$ amine (DIPEA: N,N-Diisopropylethylamine, ACN: Acetonitrile, RT: Room temperature). (**b**) Characterization of the DM1-linker by $^1$H NMR and (**c**) $^{13}$C NMR.

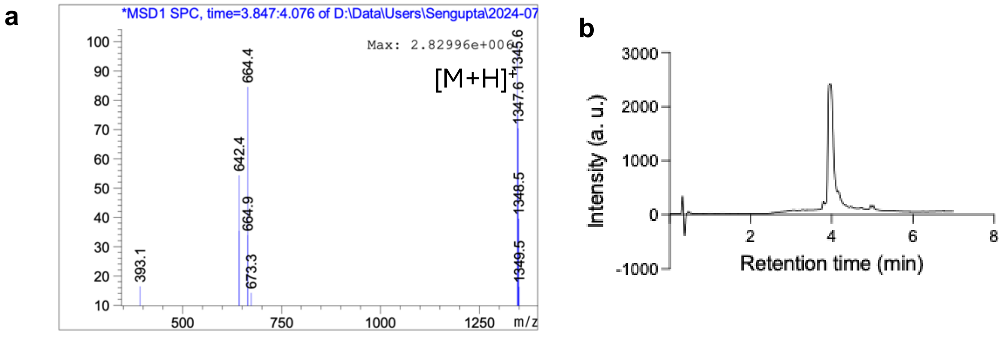


**Supplementary Figure 3:** (**a**) Mass spectral analysis of DM1 linker. Calculated mass for [M+H]+: 1345.6 Da, observed mass: 1345.6 Da. (**b**) RP-HPLC trace of DM1-linker representing the purity. A gradient method of 100 % water to 100 % acetonitrile in the presence of 0.1 % formic acid was used for HPLC using a C18 column

**a**

| Condition | Antibody : linker | Concentration of antibody | Buffer | Enzyme quantity |
|---|---|---|---|---|
| **a** | 1:50 | 2 mg/ml | 1:1 PBS+HEPES | 2.5 eq |
| **b** | 1:25 | 5 mg/ml | PBS | 2.5 eq |
| **c** | 1:50 | 2 mg/ml | 1:2.5 PBS+HEPES | 2.5 eq |
| **d** | 1:100 | 10 mg/ml | HEPES | 5 eq |
| **e** | 1:50 | 5 mg/ml | 1:1 PBS+HEPES | 2.5 eq |
| **f** | 1:50 | 5 mg/ml | 1:2 PBS+HEPES | 2.5 eq |

**b**

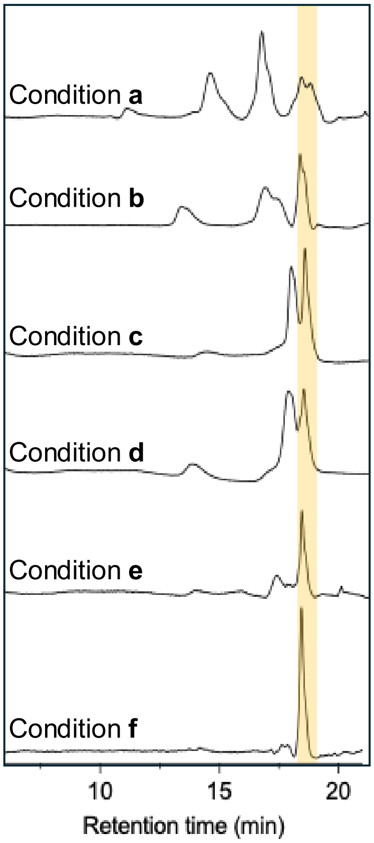


**Supplementary Figure 4: (a)** Optimization of reaction conditions for the synthesis of LT-025 using e-MTGase. **(b)** Hydrophobic interaction column chromatogram traces of the product obtained in each reaction condition. Purification using MWCO filters was carried out before running the HPLC. 10 µg of each sample was injected for analysis. Condition **f** corresponds to the chromatogram of LT-025. Method used for the HPLC is a gradient method from 100 % of 25 mM phosphate buffer (pH 7) with 1.5 M ammonium sulphate to 100 % of 25 mM phosphate buffer (pH 7) with 20 % isopropanol. The absorption was recorded at 280 nm. The chromatograms were baseline corrected using Origin 8.0 and were plotted using GraphPad Prism.

**a**

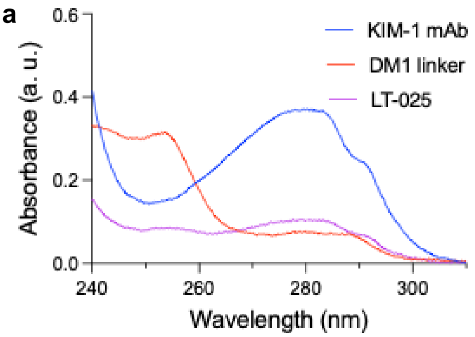


**b**

$$C_{mAb} = (A_{280}\,\varepsilon_{drug}{}^{\lambda(D)} - A_{\lambda(D)}\varepsilon_{drug}{}^{280})/[(\varepsilon_{\mathrm{mAb}}{}^{280}\,\varepsilon_{\mathrm{drug}}{}^{\lambda(\mathrm{D})} - \varepsilon_{\mathrm{mAb}}{}^{\lambda(\mathrm{D})}\,\varepsilon_{\mathrm{drug}}{}^{280})\mathrm{l}]$$

$$C_{drug} = (A_{280}\,\varepsilon_{mAb}{}^{\lambda(D)} - A_{\lambda(D)}\varepsilon_{mAb}{}^{280})/[(\varepsilon_{\mathrm{drug}}{}^{280}\,\varepsilon_{\mathrm{mAb}}{}^{\lambda(\mathrm{D})} - \varepsilon_{\mathrm{dug}}{}^{\lambda(\mathrm{D})}\,\varepsilon_{\mathrm{mAb}}{}^{280})\mathrm{l}]$$

$$DAR = Cdrug/CmAb$$

**c**

| | |
|---|---|
| Conc of antibody (M) | 1.66E-06 |
| Conc. of linker (M) | 1.70E-06 |
| Absorbance of antibody at 280 nm | 0.329 |
| Absorbance of antibody at 254 nm | 0.134 |
| Absorbance of linker at 280 nm | 0.09 |
| Absorbance of linker at 254 nm | 0.35 |
| Ext. coefficient of antibody at 280 nm ($\varepsilon_{\mathrm{mAb}}{}^{280}$) | 198192.771 |
| Ext. coefficient of antibody at 254 nm ($\varepsilon_{\mathrm{mAb}}{}^{\lambda(\mathrm{D})}$) | 80722.8916 |
| Ext. coefficient of linker at 280 nm ($\varepsilon_{\mathrm{drug}}{}^{280}$) | 1.94E+05 |
| Ext. coefficient of linker at 254nm ($\varepsilon_{\mathrm{drug}}{}^{\lambda(\mathrm{D})}$) | 2.06E+05 |
| Absorbance of ADC at 280 nm ($A_{280}$) | 0.104 |
| Absorbance of ADC at 254 nm ($A_{\lambda(\mathrm{D})}$) | 0.085 |
| Conc of antibody in ADC ($C_{mAb}$) (M) | 1.97E-07 |
| Conc of linker in ADC ($C_{drug}$) (M) | 3.36E-07 |
| DAR ($C_{drug}/CmAb$) | 1.70E+00 |

**Supplementary Figure 5:** Calculation of DAR using UV-visible spectroscopy. **(a)** The absorption of spectra of LT-025, DM1 linker, and KIM-1 antibody are given. **(b)** The equation for the calculation of DAR. **(c)** Table showing the calculation of DAR using the equation by measuring the absorbance of the antibody and linker from the ADC. The extinction coefficients of the antibody and the linker were calculated by measuring the absorbance at known concentrations of antibody and linker. The absorption maxima of the linker and antibody were obtained as 254 nm, and 280 nm, respectively. The DAR was finally calculated by the ratio of $C_{drug}$ to $C_{mAb}$. Notation ‘l’ in the equation corresponds to the path length, which was considered as 1 cm.

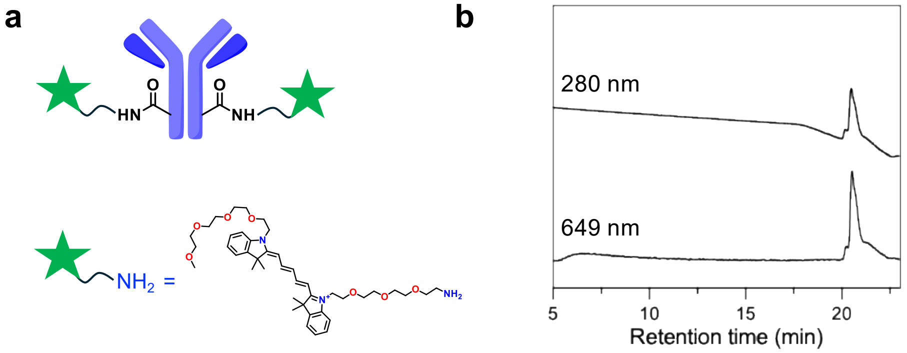


**Supplementary Figure 6: (a)** Schematic structure of KIM-1 Cy5 AFC and the structure of Cy5 linker **(b)** Characterization data of AFC using HIC. Method used for the HPLC is a gradient method from 100 % of 25 mM phosphate buffer pH 7 with 1.5 M ammonium sulphate to 100 % of 25 mM phosphate buffer pH 7 with 20 % isopropanol. The absorption was recorded at 280 nm and 649 nm. The distinguished peak in the 649 nm chromatogram represents the presence of the Cy5 fluorophore in the AFC.

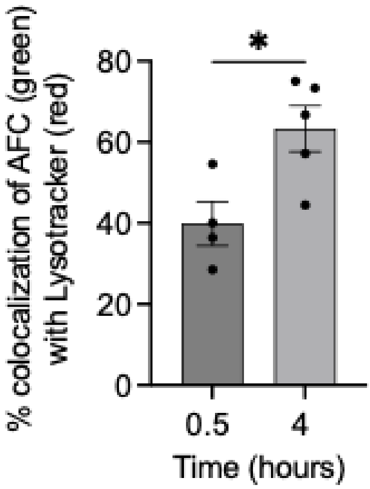


**Supplementary Figure 7:** Quantification of the colocalization between AFC (green) signal with Lysotracker (red) signal in Renca cells at 0.5 and 4 h. The data represent significantly higher colocalization at 4 h. The data were represented as mean ± SEM (n = 5, Unpaired *t*-test with Welch's correction).

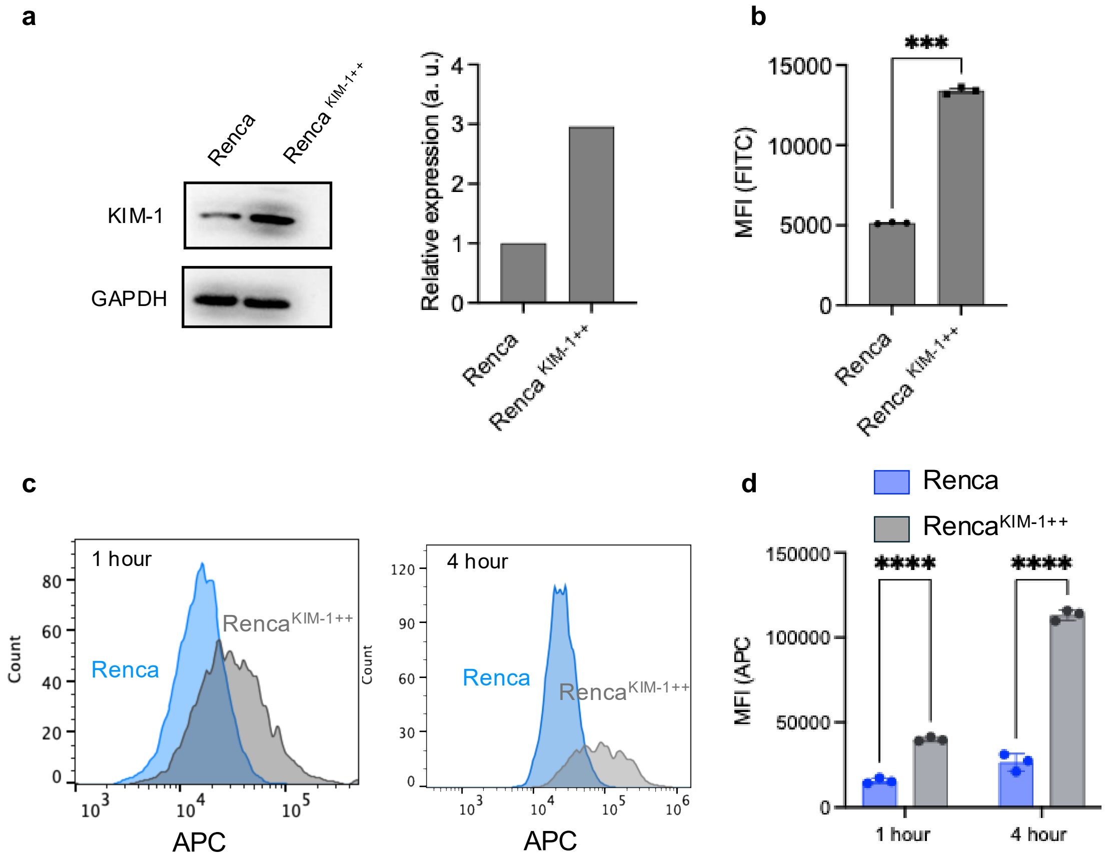


**Supplementary Figure 8:** Characterization of the KIM-1 expression on RENCA cells using (**a**) Western blotting and (**b**) flow cytometry, before and after lentiviral transduction with MR203831L3V, Origene. The mean fluorescence intensity (MFI) was plotted in the bar graph. FITC conjugated anti-mouse KIM-1 antibody was used to probe the KIM-1 expression. (**c**) The KIM-1 AFC internalization in Normal Renca and Renca$^{KIM\text{-}1++}$ by flow cytometry. The histogram shows a notable increase in the cellular internalization of AFC both at 1 and 4 h. (**d**) The bar graph represents the MFI of AFC internalization in Renca and Renca$^{KIM\text{-}1++}$ by flow cytometry. The data show a significant increase in cellular internalization. The data were represented as mean ± SEM (n = 3, two-way ANOVA).

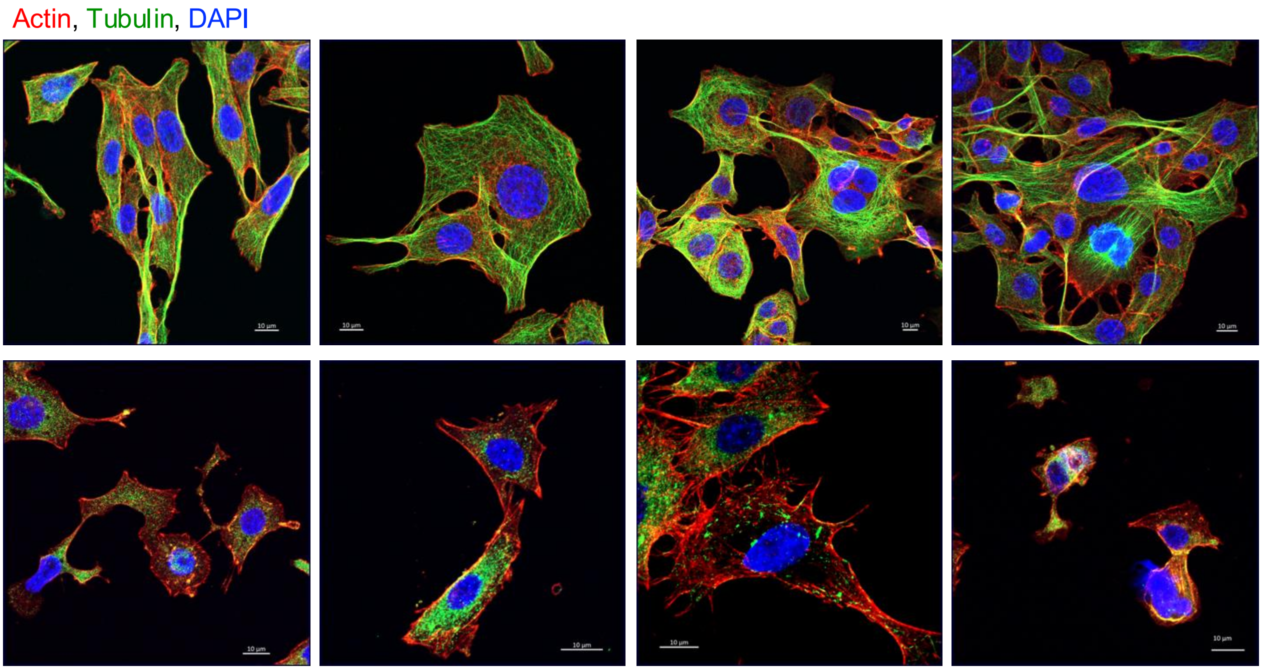


**Supplementary Figure 9:** Confocal images of Renca with and without treatment of LT-025 displaying tubulin disruption. The images showed disruption of the tubulin crosslinking (green) by DM1, a known microtubule inhibitor. Scale bar = 10 µm.

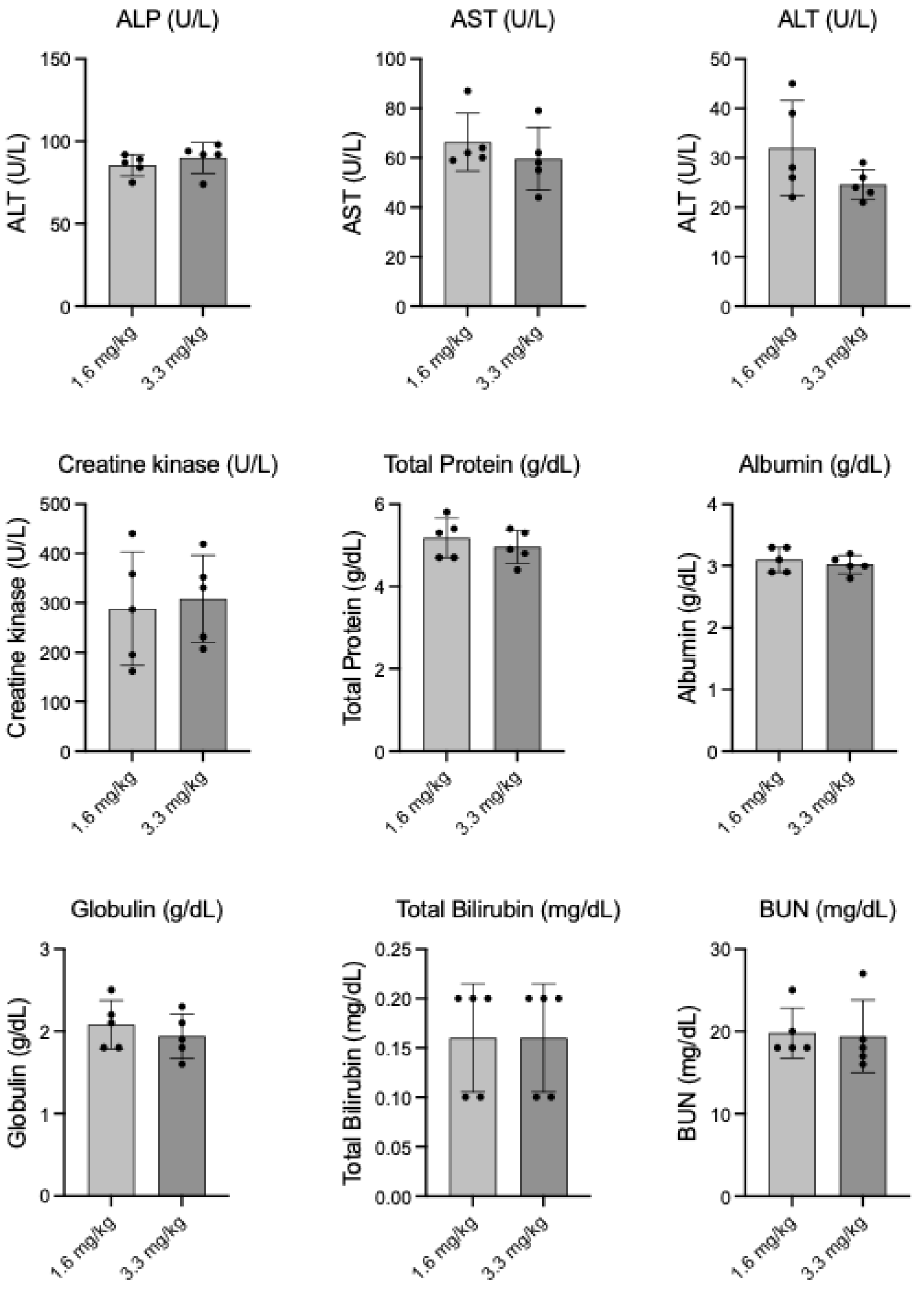


**Supplementary Figure 10:** Blood biochemistry analysis for the critical enzymes and blood components to evaluate any treatment-related toxicity in the dose escalation study by LT-025 in balb/c mice. Data shows no abnormal level of enzymes and critical blood parameters, indicating safe treatment of LT-025.

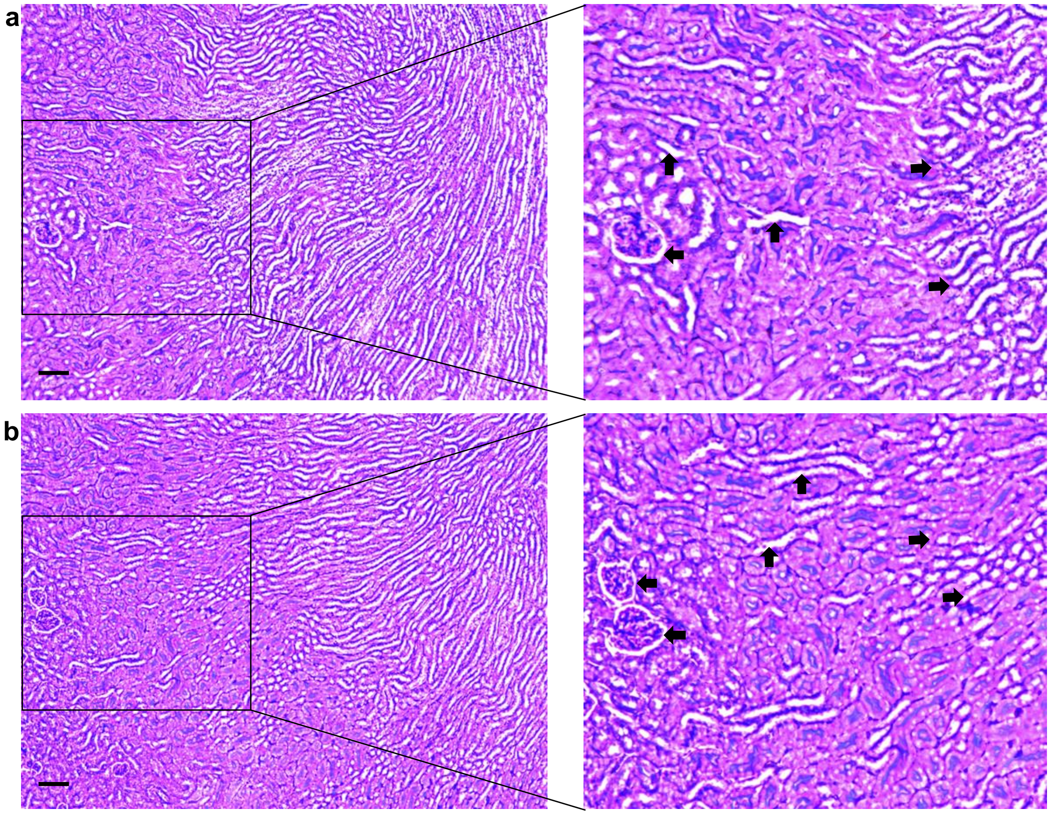


**Supplementary Figure 11:** Light microscopic images of H&E stained kidney tissue of balb/c mice in the dose escalation study by LT-025. Both the doses 1.6 mg/kg (a) and 3.3 mg/kg (b) did not show any significant renal toxicity. No signs of luminal cellular debris (up arrow), preserved Bowman's space (left arrow), and regular tubular architecture (right arrow) show no significant renal toxicity. Scale bar= 100 µm a

b

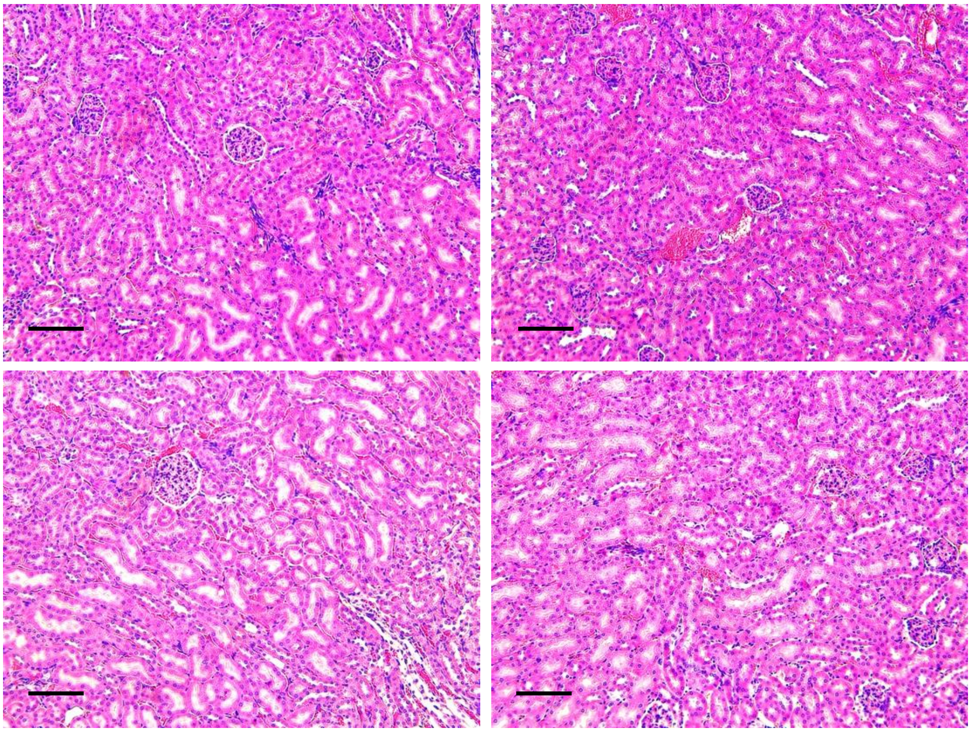

**Supplementary Figure 12:** Light microscopic images of periodic acid-Schiff (PAS) stained kidney tissue of balb/c mice in following dose escalation study with LT-025. Proximal tubules display intact PAS-positive apical brush borders and uniform tubular basement membranes, and glomerular morphology appears normal consistent with the absence of overt histopathologic nephrotoxicity under both the doses 1.6 mg/kg (a) and 3.3 mg/kg (b). Scale bar= 50 µm
